# Single-dose respiratory mucosal delivery of next-generation viral-vectored COVID-19 vaccine provides robust protection against both ancestral and variant strains of SARS-CoV-2

**DOI:** 10.1101/2021.07.16.452721

**Authors:** Sam Afkhami, Michael R. D’Agostino, Ali Zhang, Hannah D. Stacey, Art Marzok, Alisha Kang, Ramandeep Singh, Jegarubee Bavananthasivam, Gluke Ye, Xiangqian Luo, Fuan Wang, Jann C. Ang, Anna Zganiacz, Uma Sankar, Natallia Kazhdan, Joshua F. E. Koenig, Allyssa Phelps, Manel Jordana, Yonghong Wan, Karen L. Mossman, Mangalakumari Jeyanathan, Amy Gillgrass, Maria Fe C. Medina, Fiona Smaill, Brian D. Lichty, Matthew S. Miller, Zhou Xing

**Affiliations:** McMaster Immunology Research Centre, M. G. DeGroote Institute for Infectious Disease Research & Department of Medicine, McMaster University, Hamilton, ON, L8S 4K1, Canada; McMaster Immunology Research Centre, M. G. DeGroote Institute for Infectious Disease Research & Department of Biochemistry & Biomedical Sciences, McMaster University, Hamilton, ON, L8S 4K1, Canada; Department of Pediatric Otolaryngology, Shenzhen Hospital, Southern Medical University, Shenzhen, China; Department of Pathology and Molecular Medicine & M. G. DeGroote Institute for Infectious Disease Research, McMaster University, Hamilton, ON, L8S 4K1, Canada

## Abstract

The emerging SARS-CoV-2 variants of concern (VOC) increasingly threaten the effectiveness of current first-generation COVID-19 vaccines that are administered intramuscularly and are designed to only target the spike protein. There is thus a pressing need to develop next-generation vaccine strategies to provide more broad and long-lasting protection. By using adenoviral vectors (Ad) of human and chimpanzee origin, we developed Ad-vectored trivalent COVID-19 vaccines expressing Spike-1, Nucleocapsid and RdRp antigens and evaluated them following single-dose intramuscular or intranasal immunization in murine models. We show that respiratory mucosal immunization, particularly with chimpanzee Ad-vectored vaccine, is superior to intramuscular immunization in induction of the three-arm immunity, consisting of local and systemic antibody responses, mucosal tissue-resident memory T cells, and mucosal trained innate immunity. We further show that single-dose intranasal immunization provides robust protection against not only the ancestral strain of SARS-CoV-2, but also two emerging VOC, B.1.1.7 and B.1.351. Our findings indicate that single-dose respiratory mucosal delivery of an Ad-vectored multivalent vaccine represents an effective next-generation COVID-19 vaccine strategy against current and future VOC. This strategy has great potential to be used not only to boost first-generation vaccine-induced immunity but also to expand the breadth of protective T cell immunity at the respiratory mucosa.

## Introduction

Since the first report of its outbreak in Wuhan, China in late 2019, the new severe acute respiratory syndrome coronavirus 2 (SARS-CoV-2) has globally infected 187M people and claimed 4M lives and counting. Roughly 18 months into the pandemic, many countries are still battling major waves of coronavirus disease 2019 (COVID-19). In addition to general mitigation/infection control measures, the only safe and effective way to control COVID-19 is to establish herd immunity through vaccination (Fontanet and Cauchemez, 2020; Jeyanathan et al., 2020). In response to the pressing need for vaccines, a great number of candidates are being developed according to an accelerated pandemic vaccine paradigm (Lurie et al., 2020). To date, there have been at least 100 candidate vaccines tested in clinical trials and another 180 under preclinical development. These efforts have led a growing number of front-runner COVID-19 vaccines to receive emergency use authorization in various countries. Notably, several authorized first-generation vaccines are based on mRNA and recombinant adenoviral platforms designed to express the spike protein of the ancestral SARS-CoV-2 and elicit neutralizing antibody responses following 1-2 intramuscular injections (Jeyanathan et al., 2020).

The global rollout of COVID-19 vaccines has played a critical role in reducing viral transmission, hospitalizations, and deaths, particularly in a number of countries reaching high vaccine coverage rates. However, since September 2020, four SARS-CoV-2 variants of concern (VOC) have been identified and have spread rapidly across the globe, including B.1.1.7 (UK, Alpha), B.1.351 (South Africa, Beta), P.1 (Brazil, Delta) and B.1.617 (India, Gamma) (Andreano and Rappuoli, 2021; Gupta, 2021). While all of these VOC have multiple mutations in the spike protein, both B.1.351 and P.1 harbor three mutations within the RBD (K417T, E484K and N501Y) that significantly reduce their neutralization by antibodies present in either convalescent sera or COVID-19 vaccine-induced sera (Chen et al., 2021; Garcia-Beltran et al., 2021; Geers et al., 2021; Hoffmann et al., 2021b; Planas et al., 2021; Wang et al., 2021). Some B.1.617 sub-lineages also carry E484Q and L452R mutations that may reduce antibody binding (Starr et al., 2021). Of importance, several first-generation vaccines including ChAdOx1 nCoV-19 (AstraZeneca/Oxford) (Madhi et al., 2021), Ad26.COV2.S (J&J) (Sadoff et al., 2021), NVX-CoV2373 (Novavax) (Shinde et al., 2021), and BNT162b2 (Pfizer-BioNTech) (Abu-Raddad et al., 2021) have demonstrated varying degrees of reduced efficacy in protecting from mild to moderate COVID-19 caused by B.1.351. Likewise, sera from those immunized with mRNA-1273 (Moderna) have reduced neutralization potency against B.1.351 (Shen et al., 2021). Thus, there is a growing global issue that emerging VOC capable of escaping the immunity offered by first-generation vaccines may threaten to impede or disrupt the establishment and sustainability of vaccine-induced herd immunity (Aschwanden, 2021; Harvey et al., 2021).

To confront the challenges arising from VOC and uncertainty of the durability of first-generation vaccine-induced immunity, there is an urgent need to develop next-generation COVID-19 vaccine strategies (Callaway and Ledford, 2021; Gupta, 2021; Jeyanathan et al., 2020). Although updating the spike antigen to specific VOC represents one such strategy (Callaway and Ledford, 2021; Gupta, 2021), it is cumbersome and expensive, and requires selection of specific VOC sequence(s) which may result in inherently inaccurate prediction of antigenic drift, akin to current seasonal influenza virus vaccines. An alternative strategy is to develop recombinant viral-vectored multivalent vaccines amenable to respiratory mucosal immunization (Jeyanathan et al., 2020). Besides the spike antigen, such vaccines express additional conserved SARS-CoV-2 antigens to broaden T cell immunity. Since antigenic changes in conserved, internal viral proteins that are the primary focus of T cell responses are rare/improbable in SARS-CoV-2 viruses including VOC (Alter et al., 2021; Geers et al., 2021), such multivalent vaccines would not be expected to require frequent updating while they can maintain and boost the spike antigen-specific immunity induced by the first-generation vaccine. Thus, these vaccines are expected to be effective against both ancestral and variants of SARS-CoV-2. Furthermore, adenoviral vectors delivered via the respiratory mucosal route induce protection via eliciting both mucosal tissue-resident innate immune memory/trained innate immunity and adaptive immunity at the site of viral entry (Jeyanathan et al., 2020; Teijaro and Farber, 2021; Xing et al., 2020; Yao et al., 2018). Unfortunately, to-date there has been a paucity of next-generation COVID-19 vaccine strategies capable of robust and durable protection against the ancestral strain and variants of SARS-CoV-2.

In our current study, we have developed and evaluated a next-generation COVID-19 vaccine strategy in murine models. Our vaccine is built upon adenoviral vectors of human (Tri:HuAd) or chimpanzee (Tri:ChAd) origin, expressing three SARS-CoV-2 antigens (spike protein 1, full-length nucleocapsid protein, and truncated polymerase), and is suitable for respiratory mucosal delivery. We show that single-dose intranasal (i.n.) immunization, particularly with the Tri:ChAd vaccine, induces potent neutralizing antibodies both locally and systemically. Of importance, i.n. immunization elicits respiratory mucosal tissue-resident memory CD8^+^ T cells and trained resident alveolar macrophages. Such all-around mucosal immunity leads to robust protection against not only the ancestral strain of SARS-CoV-2 but also B.1.1.7 and B.1.351 variants. We further show that the i.n. route of vaccination is superior to intramuscular (i.m.) immunization at inducing protective mucosal immunity, and such protection requires both B and T cells. Our study thus indicates that respiratory mucosal delivery of a multivalent viral-vectored COVID-19 vaccine represents an effective next-generation vaccine strategy against emerging VOC.

## Results

### Construction and characterization of HuAd- and ChAd-vectored trivalent COVID-19 vaccines

Currently approved recombinant first-generation COVID-19 vaccines only encode the full-length spike (S) protein from the Wuhan-Hu-1 ancestral strain of SARS-CoV-2, and were designed primarily to induce neutralizing antibodies following intramuscular injections. This vaccine strategy is inadequate to effectively combat emerging variants of concern (VOC) (Aschwanden, 2021; Harvey et al., 2021). To address this issue, we set out to develop next-generation recombinant adenoviral-vectored COVID-19 vaccines capable of boosting the immunity induced by first-generation vaccines and broadening T cell immunity. We elected to utilize human serotype 5 (Tri:HuAd) and chimpanzee serotype 68 (Tri:ChAd) adenoviral vectors. Our vaccines were constructed to include the full-length S1 domain of spike, which in addition to numerous conserved T cell epitopes (Geers et al., 2021; Tarke et al., 2021) contains the NTD and RBD. The S1 was fused to the vesicular stomatitis virus G protein (VSVG) transmembrane/trimerization domain (**Figure 1A**) to anchor it to the membrane and facilitate its trimerization and exosomal targeting for enhanced neutralizing antibody responses (Bliss et al., 2020; Kuate et al., 2007). To broaden T cell immunity against additional viral antigens, we incorporated the full-length Nucleocapsid (N) and truncated nsp12 (RNA-dependent RNA polymerase; RdRp) proteins in our vaccine design, expressed as a single polyprotein downstream from a porcine teschovirus 2A sequence (P2A) (**Figure 1A**). N is the most abundant viral protein and is rich in immunogenic T cell epitopes in humans (Altmann and Boyton, 2020; Dai and Gao, 2021) and compared to T cell responses to S, multifunctional and universal N-specific CD8^+^ T cells are more frequently detected in convalescent COVID-19 patients (Peng et al., 2020). Furthermore, an Ad-vectored vaccine expressing N has been shown to induce protective immunity in preclinical models of SARS-CoV-2 (Class et al., 2021; Matchett et al., 2021). A region of RdRp, conserved even across other bat coronaviruses, was selected to include a maximal number of high-affinity T cell epitopes for HLA 0101, 0201, and 0301, based on bioinformatic analysis and a T cell epitope prediction tool. T cell responses to S1, N and RdRp are expected to be long-lasting and maintained following VOC infection in humans. T cells specific for N and RdRp are also cross-reactive with other bat-derived coronaviruses (Alter et al., 2021; Altmann and Boyton, 2020; Geers et al., 2021; Tarke et al., 2021).

**Figure 1.**
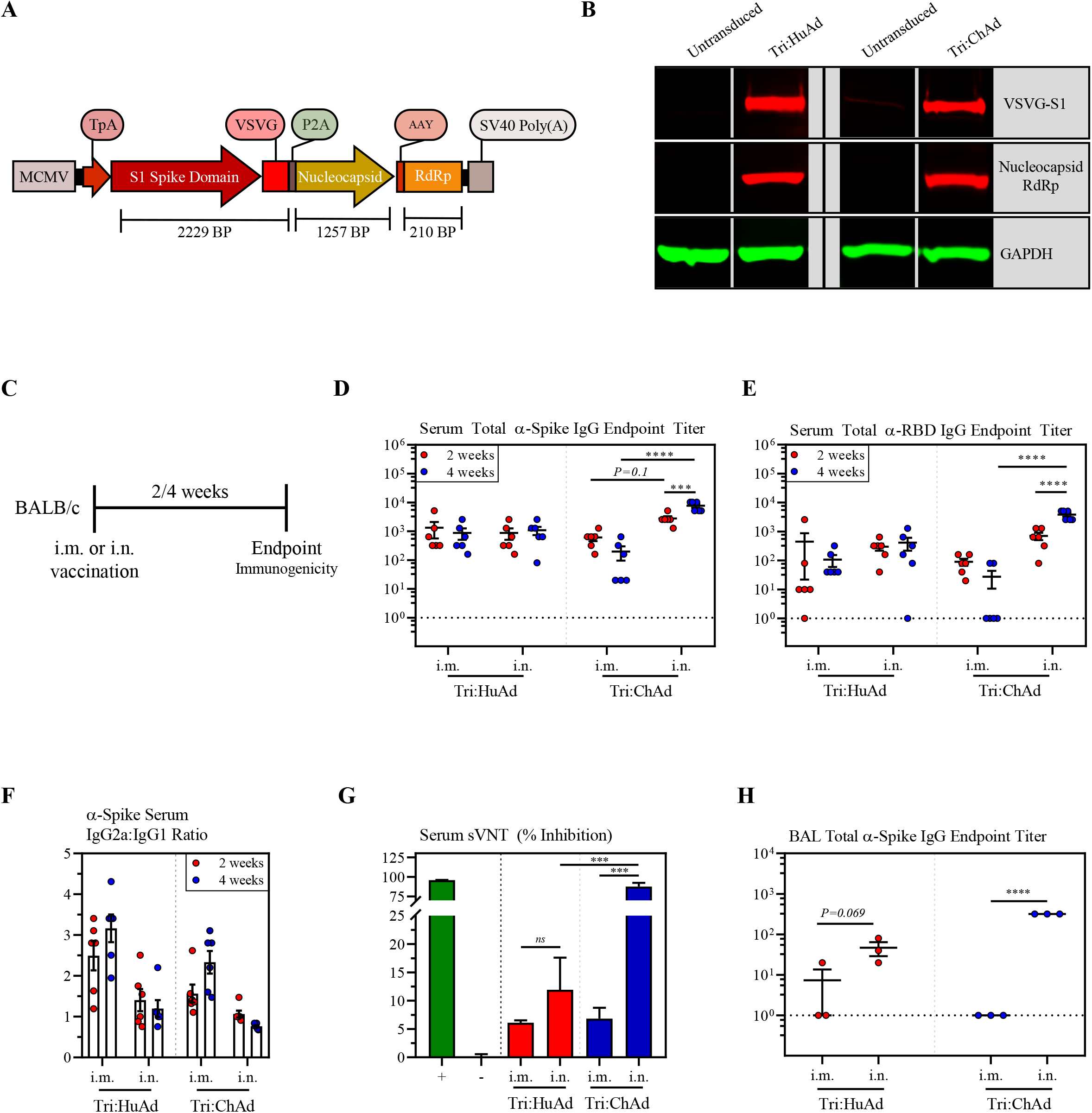
Construction and characterization of HuAd- and ChAd-vectored trivalent COVID-19 vaccines. (A) Transgene cassette diagram. The transgene construct is under control of the MCMV promoter. The S1 spike domain is fused to VSVG to facilitate trimerization and exosome targeting. Nucleocapsid and RdRp are expressed as a polyprotein, independent of S1 through a P2A site. (B) Western blot analysis of expression S1-VSVG and nucleocapsid-RdRp from whole-cell lysates from A549 cells either untransduced or infected with either Tri:HuAd or Tri:ChAd. GAPDH was used as a loading control for each condition. (C) Schema of vaccination regimen. BALB/c mice were either intramuscularly (i.m.) or intranasally (i.n.) vaccinated with a single dose of either Tri:HuAd or Tri:ChAd. A subset of animals was sacrificed 2- or 4-weeks post-immunization for immunological analysis. (D) Serum anti-spike IgG reciprocal endpoint antibody titers at 2 (red) and 4 (blue) weeks post-immunization. (E) Serum anti-RBD IgG reciprocal endpoint antibody titers at 2 (red) and 4 (blue) weeks post-immunization. (F) Reciprocal endpoint titer ratios of anti-spike IgG2a:IgG1 at 2 (red) and 4 blue) weeks post-immunization. (G) Bar graph depicting serum neutralizing antibody responses 4 weeks-post immuniztion, measured by percent (%) inhibition with a surrogate virus neutralization test (sVNT). Green bar (+) indicates assay positive control. Gray bar (-) indicates assay negative control. (H) BAL anti-spike IgG reciprocal endpoint antibody titers at 4 weeks post-immunization. Data presented in C–H represent mean ± SEM. Statistical analysis for panels D,E,H were 2-way ANOVA’s with Tukey multiple comparisons test. Statistical analysis for panel G were two-tailed T-tests. Data is representative of 1-to-2 independent experiments, n=3-6 mice/group. *p < 0.05; **p < 0.01; ***p < 0.001; ****p < 0.0001.

Prior to viral rescue, the transgene cassette was verified to be in-frame by Sanger sequencing, with translation initiating at the TpA signal sequence. Following virus rescue, amplification and purification, A549 cells were transduced with or without an MOI of 100 for Tri:HuAd and 50 for Tri:ChAd. Cells were harvested 18 hours post-infection and transgene expression at the protein level was verified by Western blot analysis for S1-VSVG, and Nucleocapsid:RdRp fusion proteins of expected sizes (**Figure 1B**). These data indicate that the transgene is in-frame, and the recombinant vaccine antigens are translated and detectable upon transduction with these two COVID-19 vaccines.

In anticipation of their further clinical evaluation in humans, we assessed the safety of Tri:HuAd and Tri:ChAd during the acute phase (within 3-days) following intramuscular (i.m.) or intranasal (i.n.) inoculation in mice (**Figure S1A**). As expected, there was little change in body weight following single-dose i.m. or i.n. vaccination (**Figure S1B**). Regardless of immunization route or vaccine vector, we observed no significant elevations in the lung or bronchoalveolar lavage fluid (BAL) in either neutrophils as assessed by FACS (**Figure S1C**) or pro-inflammatory cytokines as assessed by a multiplex-based assay (**Figure S1D**), relative to naïve controls. In accordance with these observations, serum chemistry analysis showed no indications of hepatic or renal toxicity based on alkaline phosphatase/alanine aminotransferase, and creatinine, respectively (**Figure S1E**). These data support an overall satisfactory safety profile of Tri:HuAd and Tri:ChAd COVID-19 vaccines delivered via intramuscular or respiratory mucosal route in murine models.

### Single-dose intranasal immunization leads to superior anti-spike protein humoral responses compared to intramuscular immunization

Given the close relationship between humoral responses and protective immunity in both seropositive convalescent patients and recipients of first-generation COVID-19 vaccines (Huang et al., 2020; Khoury et al., 2021; Krammer, 2021), we first examined the kinetics of spike-specific humoral immune responses following i.m. or i.n. immunization with a single-dose of either Tri:HuAd or Tri:ChAd COVID-19 vaccine. To this end, BALB/c mice were immunized and serum and BAL fluids were collected 2- and 4-weeks post-immunization (**Figure 1C**). IgG responses against spike and RBD were quantified by ELISA. Serological analysis shows that the magnitude of spike- and RBD-specific IgG responses for Tri:HuAd were similar irrespective of vaccination route, and were sustained for at least 4-weeks (**Figure 1D/E**). In contrast, i.n. Tri:ChAd vaccination induced significantly greater spike- and RBD-specific IgG responses than i.m. administration. Of note, the magnitude of these responses significantly increased from 2- to 4-weeks, an observation not seen following intramuscular immunization (**Figure 1D/E**).

Since antibody-dependent enhancement of disease (ADE) or vaccine-associated enhanced respiratory disease (VAERD) is potentially associated with Th2-biased immune responses elicited following certain viral infection and has also been experimentally observed with inactivated vaccines against SARS-CoV-1 (Bournazos et al., 2020; Jeyanathan et al., 2020), we determined the ratio of S-specific IgG2a/IgG1 antibodies as a surrogate of the Th1/Th2 immune response elicited following vaccination. Regardless of immunization route or vaccine, no Th2-skewing of humoral immune responses was seen at either assessed timepoint (**Figure 1F**).

We next assessed the neutralizing capacity of serum antibodies 4-weeks post-immunization by utilizing a commercially available surrogate Virus Neutralization test (sVNT) that allows for semi-quantitative analysis of SARS-CoV-2 neutralizing antibodies (Tan et al., 2020). A clear dichotomy in vaccine route and vector-dependent neutralizing capacity was observed, similar to those observed by ELISA (**Figure 1D/F**). Whereas immunization route had no significant effect on the neutralizing potential of serum antibodies in Tri:HuAd-vaccinated animals (i.m.: 6.1%±0.2% vs. i.n. 11.92%±2.7%), i.n. immunization with Tri:ChAd generated antibody responses with markedly enhanced neutralizing potential (87.70%±2.3%) over that by i.m. immunization (**Figure 1G**).

To assess humoral responses situated at the respiratory mucosa, BAL fluids were collected 4-weeks post-immunization with either trivalent COVID-19 vaccine and assessed for S-specific IgG responses. As expected, we were only able to reliably detect S-specific antibodies situated in the airway following i.n., but not i.m., immunization (**Figure 1H**). Of note, airway S-specific IgG responses following Tri:ChAd immunization were nearly double that of Tri:HuAd, further highlighting the superior immunogenicity of Tri:ChAd.

We next assessed the durability of antibody responses at a protracted time point, 8-weeks post-vaccination (**Figure 2A**). Interestingly, relative to 4-weeks post-immunization (**Figure 1D/E**), serum S- and RBD-specific IgG responses contracted following i.m. immunization, whereas IgG titers were sustained following i.n. immunization with either vaccine (**Figure 2B**). Once again, the serum neutralization profile determined by sVNT at 8-weeks was similar to that observed at 4-weeks post-immunization (**Figure 1G**), showing i.n. Tri:ChAd immunization to induce the highest titers of neutralizing antibodies (**Figure 2C**). Given the robust neutralizing capacity exhibited by serum from mice vaccinated intranasally with Tri:ChAd, we next tested it in a stringent microneutralization assay with live SARS-CoV-2. Congruent with the sVNT results, single-dose i.m. immunization with either vaccine afforded minimal neutralization against live SARS-CoV-2 (**Figure 2D**). In contrast, while i.n. immunization with either vaccine increased their respective neutralization capacities, Tri:ChAd vaccine elicited superior neutralization capacity over its Tri:HuAd counterpart (**Figure 2D**). As seen at 4-weeks post-immunization (**Figure 1H**), anti-S IgG from the BAL fluid was sustained at 8-weeks following i.n. immunization, more prominently with Tri:ChAd vaccine, while i.m. immunization with either vaccine failed to induce detectable anti-S IgG in the airway (**Figure 2E**).

**Figure 2.**
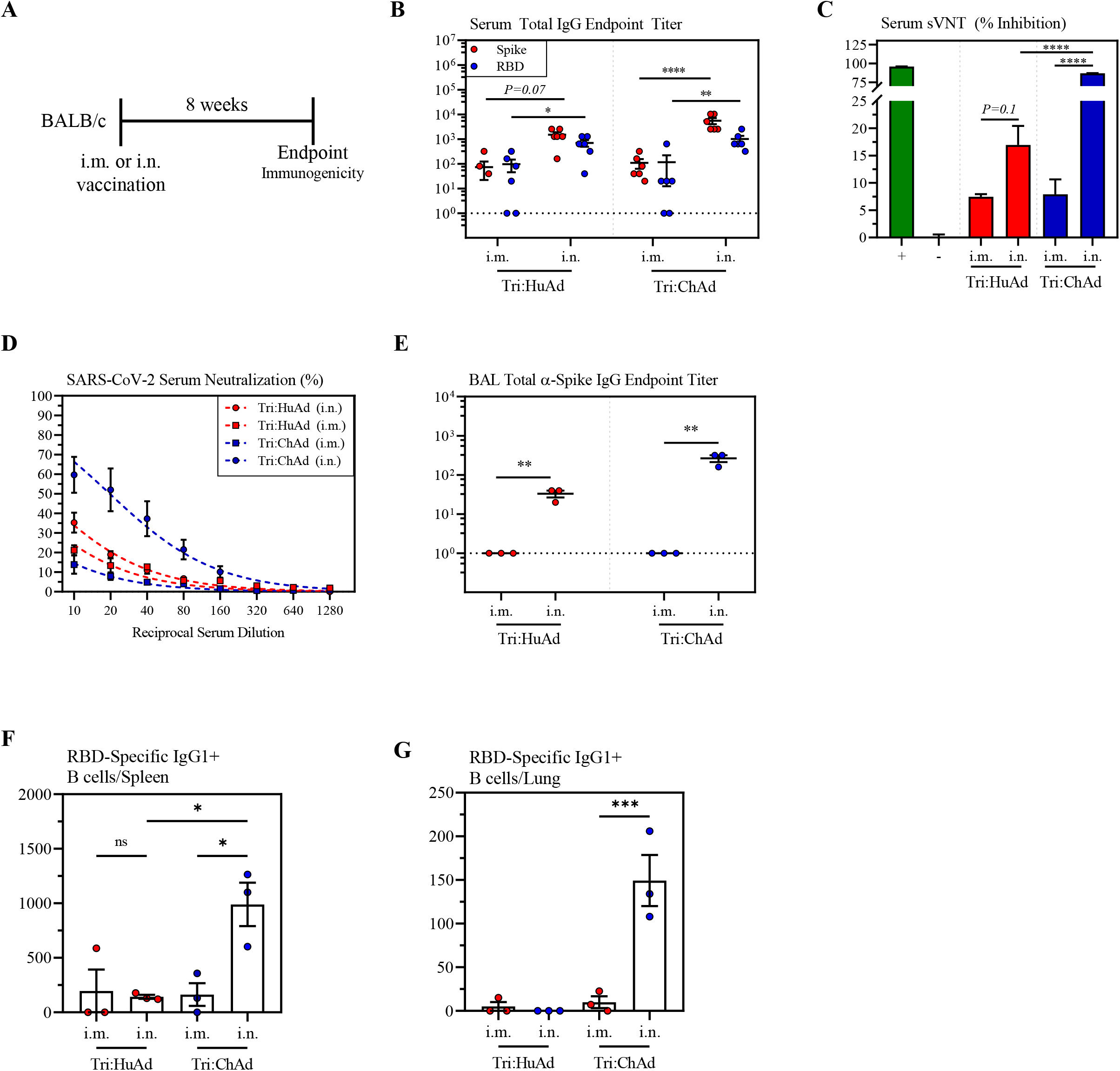
Single-dose intranasal immunization leads to superior anti-spike protein humoral responses over intramuscular immunization. (A) Schema of vaccination regimen. BALB/c mice were either i.m. or i.n. vaccinated with a single dose of either Tri:HuAd or Tri:ChAd. Animals were sacrificed 8 weeks post-immunization for immunological analysis. (B) Serum anti-spike (red) or anti-RBD (blue) IgG reciprocal endpoint antibody titers at 8 weeks post-immunization. (C) Bar graph depicting serum neutralizing antibody responses 8 weeks-post vaccination, measured by percent (%) inhibition with a surrogate virus neutralization test (sVNT). Green bar (+) indicates assay positive control. Gray bar (-) indicates assay negative control. Assay run in parallel to that in Figure 1G, utilizing the same positive and negative plate controls. (D) Serum neutralizing antibody responses at 8 weeks post-immunization, measured by percent (%) neutralization utilizing a live SARS-CoV-2 microneutralization (MNT) assay. (E) BAL anti-spike IgG reciprocal endpoint antibody titers at 8 weeks post-immunization. (F) Bar graph depicting the absolute number of class-switched IgG1^+^ RBD-specific B cells in the spleen at 8 weeks post-immunization. (G) Bar graph depicting the absolute number of class-switched IgG1^+^ RBD-specific B cells in the lung at 8 weeks post-immunization. Data presented in B–G represent mean ± SEM. Statistical analysis for panels B, D-G were 2-way ANOVA’s with Tukey multiple comparisons test. Statistical analysis for panel C were two-tailed T-tests. Data is representative of 2 independent experiments, n=3-6 mice/group. *p < 0.05; **p < 0.01; ***p < 0.001; ****p < 0.0001.

To examine the relationship of vaccine vector and immunization route to detectable antigen-experienced memory B cells present in systemic lymphoid and local lung tissues, we tetramerized biotinylated RBD conjugated to a fluorochrome, and probed for RBD-specific B cells by flow cytometry (Hartley et al., 2020; Rodda et al., 2021). A decoy tetramer was included during staining to remove construct-specific B cells, ensuring any cells detected were bona fide RBD-specific B cells. Remarkably, while all immunizations induced a detectable population of RBD-specific B cells in the spleen, i.n. Tri:ChAd induced significantly higher levels than i.n. Tri:HuAd immunization (**Figure 2F**). In addition, only i.n. immunization with Tri:ChAd induced a distinct detectable population of RBD-specific B cells in the lung tissue (**Figure 2G**).

Together, the above data indicate that single-dose intranasal immunization, particularly with Tri-ChAd vaccine, induces superior functional humoral responses over the intramuscular route of immunization.

### Single-dose intranasal immunization induces superior airway T cell responses over intramuscular immunization

Having established the profile of vaccine-induced humoral immune responses, we next examined T cell responses with a focus on those within the airways. Besides antibodies, airway T cells are on the frontline of host defense, playing pivotal roles in immunity against emerging human coronaviruses, including SARS-CoV-2 (Jeyanathan et al., 2020; Zhao et al., 2016). To this end, BALB/c mice were immunized intramuscularly or intranasally with a single-dose of either trivalent COVID-19 vaccine, and mononuclear cells from the BAL fluid were harvested 2- and 4-weeks post-immunization. Antigen-specific T cells were analyzed by flow cytometry for intracellular IFNγ, TNFα, IL-2, and Granzyme B expression upon *ex vivo* stimulation with 15mer peptide pools for S1 (132 peptides), N (82 peptides), and RdRp (12 peptides). In agreement with our previously published work (Jeyanathan et al., 2017; Lai et al., 2015; Santosuosso et al., 2005), i.m. immunization failed to induce detectable antigen-specific CD8^+^ T cells in the airways, irrespective of vaccine vector (**Figure 3A/D/G**). In stark contrast, i.n. immunization induced a significant number of IFNγ^+^CD8^+^ T cells specific for S1, N, or RdRP in the airways. Of interest, S1-specific T cell responses were dominant relative to those for N or RdRP. Compared to Tri:HuAd, i.n. Tri:ChAd vaccine induced greater levels of airway CD8^+^ T cell responses to the three antigens, particularly at the 2-week timepoint (**Figure 3A/D/G**). Intranasal, but not intramuscular, immunization also induced a similar profile of antigen-specific CD4^+^ T cell responses in the airways, but to a lesser degree than the CD8^+^ T cell response (**Figure S2A**).

**Figure 3.**
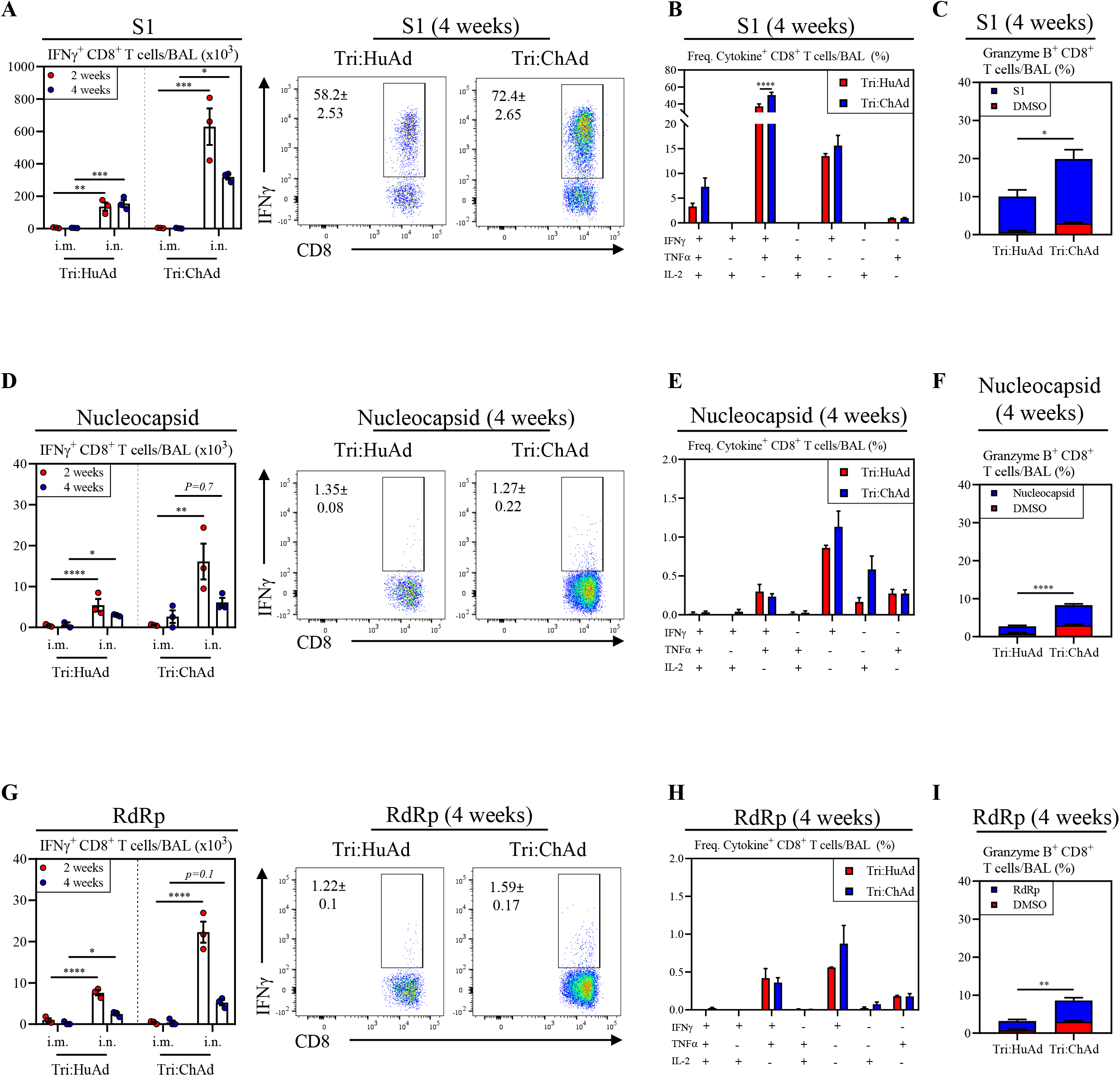
Single-dose intranasal immunization induces superior airway T cell responses over intramuscular immunization. (A) Left panel. Bar graphs depicting absolute number of S1-specific IFNɤ^+^CD8^+^ T cells in the BAL at 2 (red) and 4 (blue) weeks post-immunization (as in Figure 1C). Right panel. Representative flow cytometric dot plots of IFNɤ^+^CD8^+^ T cells in the BAL following *ex vivo* stimulation with overlapping peptide pools for S1. (B) Bar graphs depicting multifunctional CD8^+^ T cell responses in the BAL as measured by production of IFNγ, TNFα, and/or IL-2, at 4 weeks post-immunization, following *ex vivo* stimulation with overlapping peptide pools for S1. (C) Stacked bar graph depicting the frequency of cytotoxic CD8^+^ T cells in the BAL as measured by Granzyme B production at 4 weeks post-immunization, following *ex vivo* stimulation with DMSO (red) or overlapping peptide pools for S1 (blue). (D) – (F) is the same as (A) – (C) but following stimulation with overlapping peptide pools for nucleocapsid. (G) – (I) is the same as (A) – (C) but following stimulation with overlapping peptide pools for RdRp. Data presented in B–G represent mean ± SEM. Statistical analysis for panels A, B, D, E, G, H were 2-way ANOVA’s with Tukey multiple comparisons test. Statistical analysis for panel C, F, and I were two-tailed T-tests. Data is representative of 1-to-2 independent experiments, n=3-6 mice/group. *p < 0.05; **p < 0.01; ***p < 0.001; ****p < 0.0001.

We further assessed the multifunctionality of CD8^+^ T cells specific for S1, N, and RdRp following intranasal immunization. We focused on examining their IFNγ, TNFα, IL-2 and granzyme B production 4-weeks post-i.n. immunization when these cells entered the contraction/memory phase. We found that the majority of S1-specific CD8^+^ T cells induced by either Tri:HuAd or Tri:ChAd vaccine were multifunctional, co-expressing IFNγ and TNFα, whereas N- and RdRp-specific T cells were pre-dominantly monofunctional (**Figure 3B/E/H**). We further examined granzyme B production as an indicator of their cytotoxic capability. While i.n. immunization with both vaccines generated cytotoxic CD8^+^ T cells against all vaccine-encoded antigens, Tri:ChAd vaccine induced significantly greater frequencies of these T cells in the airways than Tri:HuAd (**Figure 3C/F/I**).

To compare the local respiratory mucosal responses to those induced at peripheral sites, we also assessed antigen-specific T cell responses in the spleen at 4-weeks post-immunization. In keeping with our previous findings (Santosuosso et al., 2005), contrasting what was observed at the respiratory mucosa, intramuscular, but not intranasal, immunization with either vaccine induced robust systemic CD8^+^ T cell responses to S1, N, and RdRp (**Figure S2B, top panels**). Once again, S1-specific T cell responses were immunodominant to those induced by N or RdRp. Antigen-specific CD4^+^ T cells were also induced in the spleen, but again to a lesser degree than the CD8^+^ T cell response (**Figure S2B, bottom panels**).

Collectively, the above findings indicate that single-dose intranasal, but not intramuscular, immunization, particularly with Tri:ChAd vaccine, is able to induce multifunctional CD8^+^ T cells with cytotoxic potential within the respiratory tract.

### Single-dose intranasal, but not intramuscular, immunization induces multifunctional respiratory mucosal tissue-resident memory T cell responses

Compelling evidence indicates a critical role of mucosal tissue-resident memory T cells (T_RM_) in host defense (Szabo et al., 2019) and yet induction of long-lasting lung mucosal T_RM_ is highly dependent on vaccination route. Respiratory mucosal, but not intramuscular, route of vaccination represents a robust way to induce such T cells (Jeyanathan et al., 2018, 2020; Teijaro and Farber, 2021). To investigate whether immunization with our Ad-vectored trivalent COVID-19 vaccines was able to induce persisting respiratory mucosal T_RM_, we first established t-SNE maps based on pooled CD3^+^/CD8^+^/CD4^-^ mononuclear cells isolated from whole lung tissues of all animals 8-weeks post- i.m. or i.n. immunization with either Tri:HuAd or Tri:ChAd vaccines (**Figure 4A, top panel**). Upon overlaying these cell populations concatenated from i.m. and i.n. animals, two unique CD8^+^ T cell clusters (yellow color) were identified to be associated only with i.n. route of immunization (**Figure 4A, bottom panel**). We next generated heatmaps to overlay expression intensities for the surface markers CD44, CD69, CD103 and CD49a often associated with mucosal T_RM_ (Szabo et al., 2019). This analysis shows the two unique clusters of T cells of respiratory mucosal CD8^+^ T_RM_ phenotype induced by i.n. immunization (**Figure 4B/C**). We further quantified absolute numbers of CD8^+^ T_RM_ in whole lung tissues according to immunization routes and vaccine vectors. While i.n., but not i.m., immunization with either Tri:HuAd or Tri:ChAd vaccine induced significant numbers of CD8^+^ T_RM_, Tri:ChAd vaccine induced profoundly more CD8^+^ T_RM_ in whole lung tissues (**Figure 4D, Figure S3A**).

**Figure 4.**
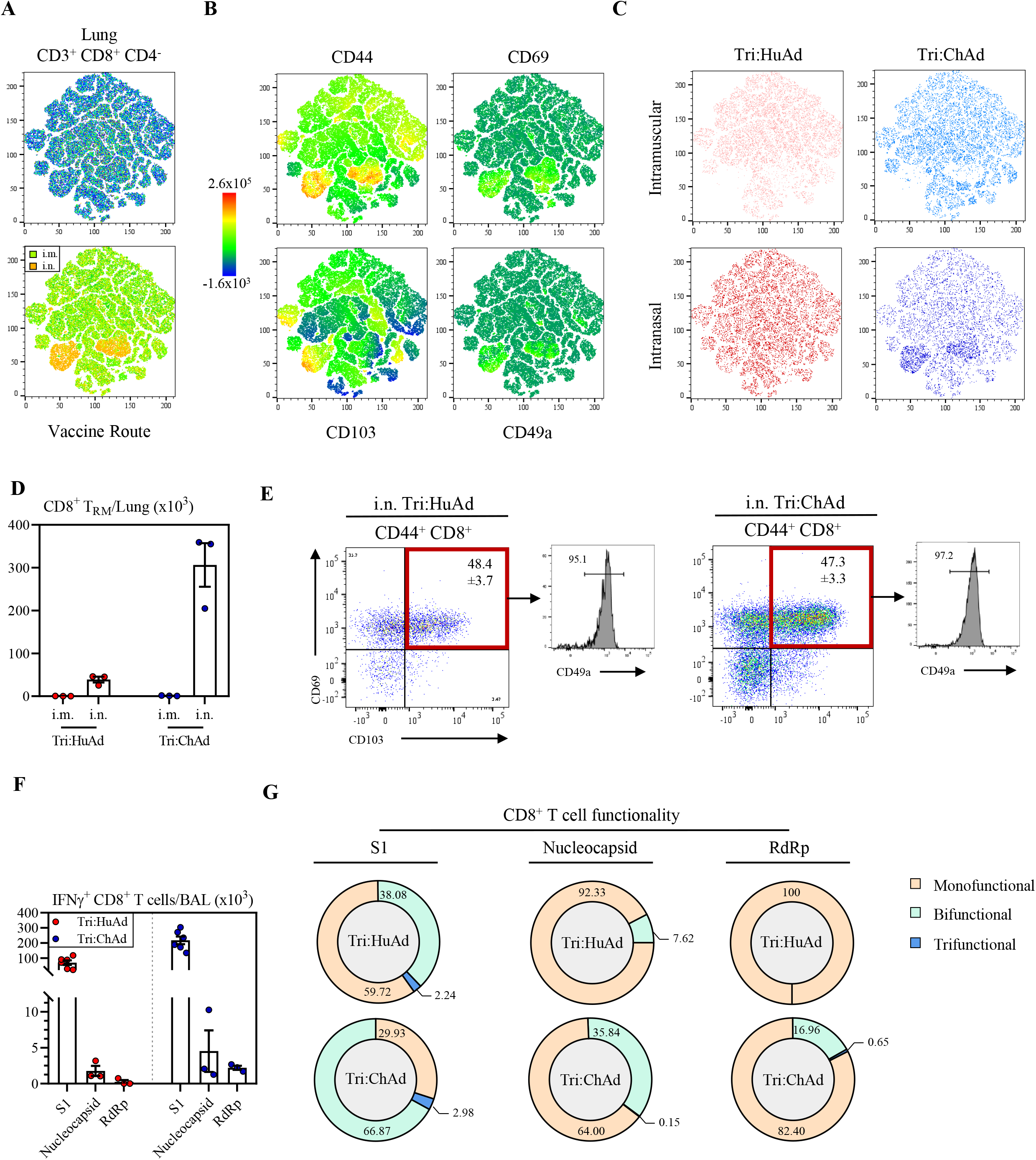
Single-dose intranasal, but not intramuscular, immunization induces multifunctional respiratory mucosal tissue-resident memory T cells. (A) Top panel: t-SNE maps were generated from concatenating CD3^+^ CD8^+^CD4^-^ gated lung mononuclear cells from 12 individual animals (3 per group of route/vaccine). Analysis was performed utilizing default FlowJo V.10 software settings. Bottom panel: Overlay of populations arising after intramuscular (green) or intranasal (yellow) onto t-SNE maps. (B) Heatmap projections of CD44, CD69, CD103, or CD49a on t-SNE maps. (C) Overlap of populations arising after intramuscular or intranasal Tri:HuAd (red) or intramuscular or intranasal Tri:ChAd (blue). (D) Bar graph depicting absolute number of tissue-resident memory CD8^+^ T cells in the lung at 8 weeks post-immunization. (E) Left panel: Flow cytometric dot plots of CD44^+^CD8^+^ T cells for CD69 and CD103 from the BAL at 8 weeks post-immunization. Right panel: Histogram depicting expression of CD49a on CD69/CD103 double-positive CD44^+^CD8^+^ T cells. (F) Bar graphs depicting absolute number of S1, nucleocapsid, or RdRp-specific IFNγ^+^ CD8^+^ T cells in the BAL at 8 weeks post-immunization. (G) Sunburst plots depicting functionality of CD8^+^ T cells at 8 weeks post-immunization, following *ex vivo* stimulation with either S1, nucleocapsid, or RdRp peptide pools. Data is representative of 1-to-2 independent experiments, n=3-6 mice/group.

We next examined induction of CD8^+^ T_RM_ within the airways (BAL) 8-weeks post-immunization. Given the lack of airway T cell responses at earlier timepoints following i.m. immunization (**Figure 3A/D/G**), we focused our analysis on BAL cells from i.n.-immunized animals. We found that regardless of the vaccine vector, approximately 50% of antigen-experienced CD44^+^CD8^+^ T cells in the BAL co-expressed T_RM_ surface markers CD69, CD103, and CD49a (**Figure 4E**). Similar observations were made with antigen-experienced CD69^+^CD11a^+^ CD4^+^ T_RM_ in the airways except that they were present in smaller frequencies compared to CD8^+^ T_RM_ (**Figure S3B**).

We further determined the multi-functionality of long-term antigen-specific memory CD8^+^ T cells in the airways 8-weeks post-i.n. immunization. Compared to the 4-week timepoint (**Figure 3A/D/G**), airway antigen-specific IFNγ^+^CD8^+^ T cell populations further contracted, irrespective of vaccine vector, with the majority being specific for S1 (**Figure 4F**). However, multi-cytokine expression analysis reveals that the antigen-specific memory CD8^+^ T cells induced by i.n. Tri:ChAd immunization were more functional than those induced by Tri:HuAd, vaccine based on the co-expression of IFNγ, TNFα, and/or IL-2 (**Figure 4G**). Of note, S1-specific memory T cells showed a greater breadth of multifunctionality than those specific to N or RdRP.

Altogether, these data indicate that intranasal, but not intramuscular, route of COVID-19 immunization is able to induce durable multifunctional respiratory mucosal T_RM_ responses. Furthermore, Tri:ChAd vaccine is more potent than Tri:HuAd platform.

### Single-dose intranasal, but not intramuscular, immunization induces trained airway macrophages

Since alveolar macrophages (AM) are the most prominent tissue-resident phagocyte stationed on the surface of the respiratory mucosa and have been shown to interact with SARS-CoV-2 (Grant et al., 2021) and other RNA viruses (Kumagai et al., 2007; Schneider et al., 2014), they likely play a critical role in early innate immune control of SARS-CoV-2 infection. Aside from the induction of robust adaptive immunity within the airways, we have recently shown that Ad-vectored respiratory mucosal, but not intramuscular, TB immunization is capable of generating long-lasting airway-resident memory AM and trained innate immunity (D’Agostino et al., 2020; Xing et al., 2020; Yao et al., 2018). Such memory or trained AM are phenotypically defined by their markedly increased surface expression of MHC II (Yao et al., 2018). Thus, to investigate whether immunization with Ad-vectored COVID-19 vaccines induces trained AM, mice were immunized intramuscularly or intranasally with Tri:HuAd or Tri:ChAd vaccine and the immune phenotype of airway (BAL) macrophages was analyzed 8-weeks post-immunization. (**Figure 5A**). To better enable our flow cytometric characterization of CD45^+^CD11b^+^CD11c^+^ airway macrophages, t-SNE maps were first generated with the BAL cells pooled from all of animals to visualize the overall differences between both vaccine vectors and routes (**Figure 5B/C**). AM were easily distinguishable as a large cluster co-expressing high levels of AM surface markers CD11c and Siglec-F (**Figure 5B)**. Of importance, while high MHC II expression was seen within the large AM cluster, another clear cluster branched off the main AM population, co-expressing high levels of MHC II and CD11b (**Figure 5B, top right panels**). A small neutrophil cluster was discernable by high co-expression of both CD11b and Ly6G (**Figure 5B**).

**Figure 5.**
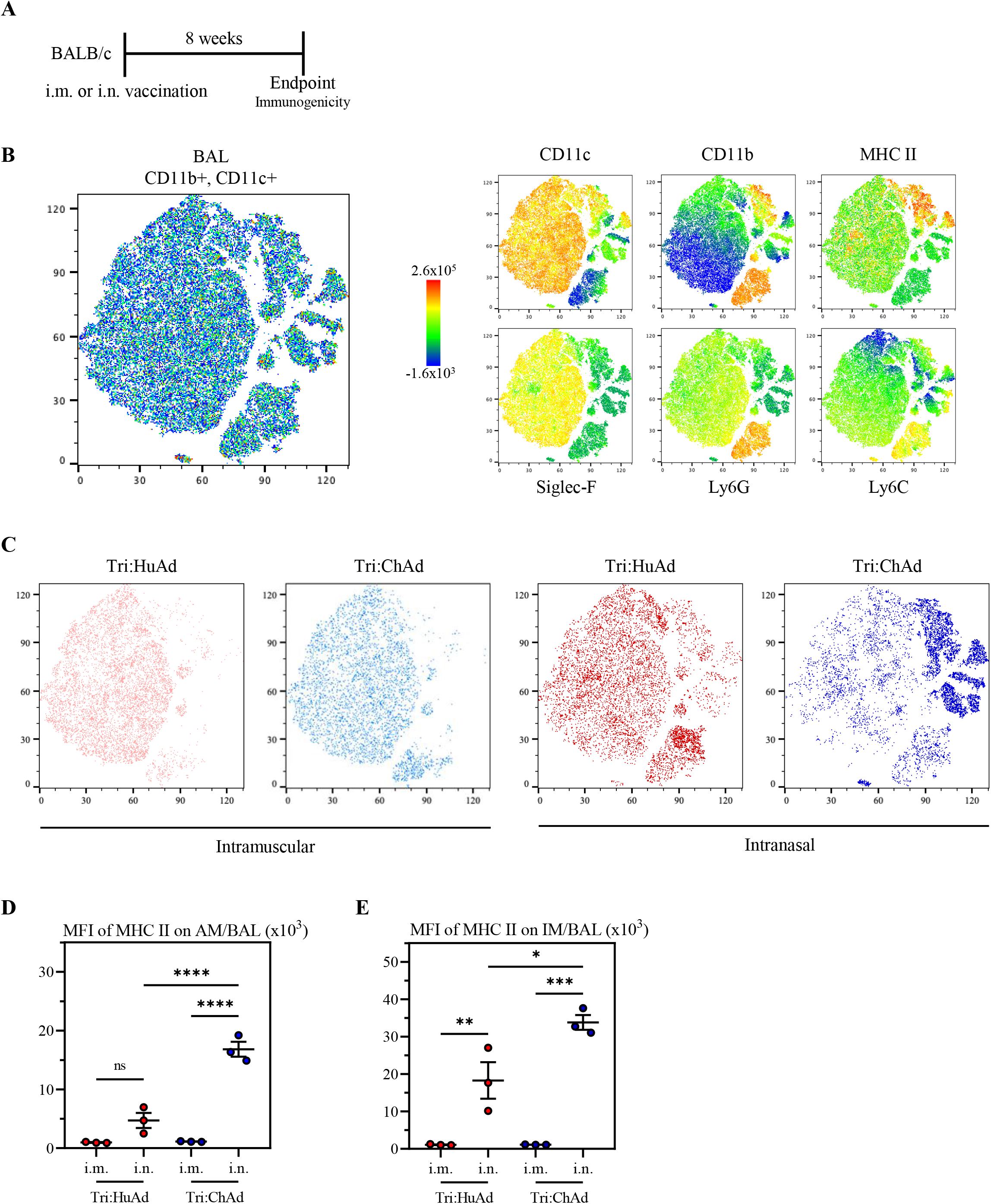
Single-dose intranasal, but not intramuscular, immunization induces trained airway macrophages. (A) Schema of vaccination regimen. BALB/c mice were i.m. or i.n. vaccinated with a single dose of either Tri:HuAd or Tri:ChAd. Animals were sacrificed at 8 weeks post-immunization for immunological analysis. (B) Left panel: t-SNE maps were generated from concatenating CD45^+^CD11b^+^CD11c^+^ gated BAL mononuclear cells from 12 individual animals (3 per group of route/vaccine condition). Analysis was performed utilizing default FlowJo V.10 software settings. Right panels: Heatmap projections of CD11c, CD11b, MHC II, Siglec-F, Ly6G or Ly6C on t-SNE maps. (C) Overlap of populations arising after intramuscular or intranasal Tri:HuAd (red) or intramuscular or intranasal Tri:ChAd (blue). (D) MFI of MHC II expression on AM in BAL at 8 weeks post-immunization. (E) MFI of MHC II expression on IM in BAL at 8 weeks post-immunization. Data presented in D–E represent mean ± SEM from n=3 mice/group. Statistical analysis for panels D, E were 1-way ANOVA with Tukey multiple comparisons test. *P < 0.05; **P < 0.01; ***P < 0.001; ****P < 0.0001.

We next sought to specifically determine the extent to which COVID-19 vaccine vectors and routes induced trained MHC II^high^ airway macrophages. Whereas intramuscular immunization with either Tri:HuAd or Tri:ChAd vaccine resulted in hardly any MHC II^high^ AM (**Figure 5C, left 2 panels**), intranasal immunization, most notably with Tri:ChAd, generated several distinct populations expressing high levels of MHC II or CD11b separate from the large AM cluster (**Figure 5C, right 2 panels**). This is phenotypically consistent with an influx of interstitial macrophages (IM), and trained AM observed within the airway following respiratory mucosal vaccination with a HuAd-vectored TB vaccine (Yao et al., 2018). Thus, using a comprehensive gating strategy, we compared MHC II expression in both AM and IM populations in the airways. While in keeping with our previous findings with HuAd-vectored TB vaccine (D’Agostino et al., 2020; Yao et al., 2018), i.m. immunization with Tri:HuAd or Tri:ChAd COVID-19 vaccine was unable to induce trained AM (**Figure 5D**) or IM (**Figure 5E**), i.n. immunization with either COVID-19 vaccine induced markedly increased MHC II^high^-expressing AM and IM (**Figure 5D/E**). Furthermore, in line with our visual observations by t-SNE analysis (**Figure 5C**), i.n. Tri:ChAd immunization induced significantly further increased MHC II MFI on both AM and IM over that by Tri:HuAd immunization (**Figure 5 D/E**). Collectively, the above data suggest that single-dose respiratory mucosal immunization with Ad-vectored COVID-19 vaccines also induces memory/trained airway macrophages, besides its potent effects on inducing memory adaptive B and T cell responses at the respiratory mucosa.

### Intranasal, but not intramuscular, immunization provides potent B and T cell-dependent protection from SARS-CoV-2 infection

Our immunological findings thus far indicate that respiratory mucosal immunization with our trivalent COVID-19 vaccines generates long-lasting, highly functional adaptive and innate immune responses situated at the portal of infection, the lungs. As such, we next sought to assess the protective efficacy of both vaccine candidates using a mouse-adapted virus (SARS-CoV-2 MA-10) (Leist et al., 2020). To this end, wildtype BALB/c mice were intramuscularly or intranasally immunized with a single-dose of Tri:HuAd or Tri:ChAd and subsequently challenged 4-weeks later with a lethal dose of 1×10^5^ PFU SARS-CoV-2 MA-10 (**Figure 6A**). Animals were monitored for weight loss and mortality. All unvaccinated animals succumbed, reaching humane endpoint by 5-days post-infection (**Figure 6B**). Likewise, single-dose i.m. immunization with either Tri:HuAd or Tri:ChAd vaccine failed to protect against this stringent infectious dose of SARS-CoV-2, with 80% animals reaching humane endpoint by 4-5 days post-infection (**Figure 6B;** numbers of surviving animals in each group indicated in brackets after legends). In contrast, animals intranasally immunized with Tri:HuAd vaccine showed slight (5%), transient weight loss but rapidly rebounded to pre-infection weights over the course of experimentation (**Figure 6B**). Notably, animals immunized intranasally with Tri:ChAd vaccine were fully protected from any morbidity, showing no weight loss throughout the course of experimentation (**Figure 6B**). To further evaluate vaccine-mediated protection, a cohort of mice was sacrificed 2-days post-infection for determination of viral burden during the acute phase of the infection within the lung. In accordance with weight loss, i.m. immunization with either vaccine only modestly reduced pulmonary viral loads (**Figure 6C**). However, in keeping with differentially improved clinical outcomes (**Figure 6B**), i.n. immunization provided vaccine vector-dependent reductions in viral burden, with Tri:ChAd-immunized animals having the most significantly reduced viral burden (>3 log) compared to >1 log reduction with Tri:HuAd immunization (**Figure 6C**). These data indicate a correlation between vaccine-induced measures of mucosal B and T cell immunity, as described earlier, with protective efficacy against acute lethal SARS-CoV-2 infection.

**Figure 6.**
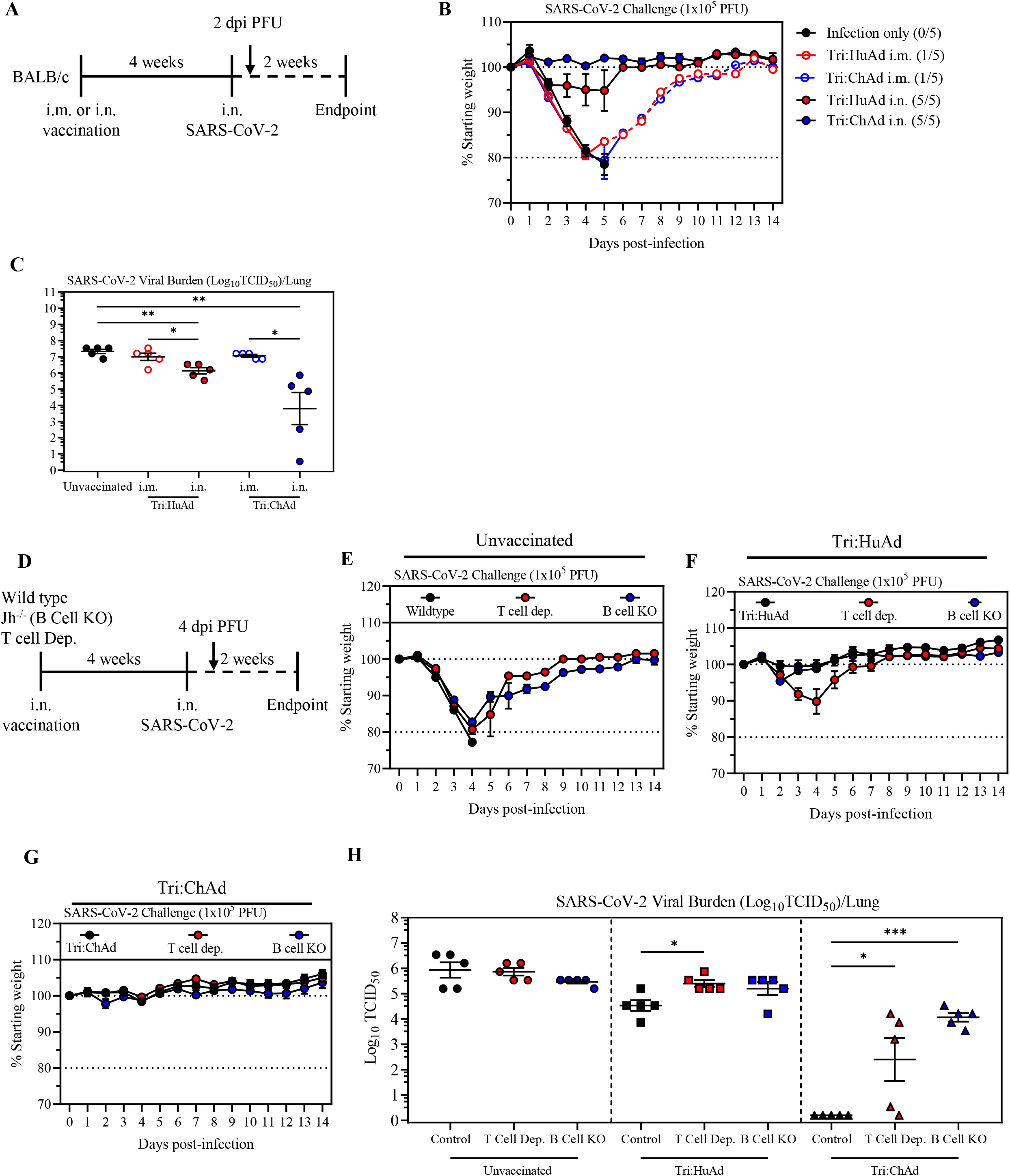
Intranasal, but not intramuscular, immunization provides potent B and T cell-dependent protection from SARS-CoV-2 infection. (A) Schema of SARS-CoV-2 challenge regimen. BALB/c mice were either i.m. or i.n. vaccinated with a single dose of either Tri:HuAd or Tri:ChAd and were infected i.n. with a lethal dose of SARS-CoV-2 MA-10, at 4 weeks post-immunization. A subset of animals was sacrificed 2 days post-infection for assessing viral burden in the lung. (B) Changes in body weight over 2 weeks post-SARS-CoV-2 infection. (C) Viral burden (Log_10_TCID_50_) in the lung at 2 days post-SARS-CoV-2 MA-10 infection. (D) Schema of SARS-CoV-2 challenge regimen. Wildtype BALB/c or Jh^-/-^ (B cell KO) mice were vaccinated i.n. with a single dose of either Tri:HuAd or Tri:ChAd for 4 weeks. A subset of BALB/c mice received intraperitoneal (i.p.) injections of CD4 and CD8 T cell depleting mAbs at both 3 and 1 days before infection. Mice were infected i.n. with a lethal dose of SARS-CoV-2 MA-10. A subset of animals was sacrificed 4 days post-infection for enumeration of viral burden. (E) Changes in body weight of unvaccinated BALB/c, T cell depleted BALB/c, or Jh ^-/-^ mice over 2 weeks post-SARS-CoV-2 infection. (F) Changes in body weight of i.n. Tri:HuAd vaccinated BALB/c, T cell depleted BALB/c, or Jh ^-/-^ mice for 2 weeks post-SARS-CoV-2 infection. (G) Changes in body weight of i.n. Tri:ChAd vaccinated BALB/c, T cell depleted BALB/c, or Jh ^-/-^ mice for 2 weeks post-SARS-CoV-2 infection. (H) Viral burden (Log_10_TCID_50_) in the lung at 4 days post-infection. Data presented in B,C,E,F,G,H represent mean ± SEM from n=5 mice/group. Statistical analysis for panels C, H were 1-way ANOVA with Tukey multiple comparisons test. *P < 0.05; **P < 0.01; ***P < 0.001.

We next investigated the relative contribution of B and T cell immunity to intranasal vaccine-induced protection. To this end, wildtype BALB/c mice (wildtype), BALB/c mice deficient in J segments (B cell KO) of the immunoglobulin heavy chain locus (therefore lacking any mature B cells or circulating IgG), and BALB/c mice depleted of CD4^+^/CD8^+^ T cells (T cell dep.) were left unvaccinated or intranasally vaccinated with Tri:HuAd vaccine. Mice were then infected with SARS-CoV-2 MA-10 at 4-weeks post-immunization (**Figure 6D**). T cell depletion was carried out from day 26 post-immunization when B cells/antibody responses were fully developed and prior to viral challenge by 2-repeated intraperitoneal injections of a T cell depleting antibody cocktail using a well-established protocol (Yao et al., 2018). Unvaccinated wildtype mice rapidly succumbed to infection, reaching humane endpoint by 4-days post-infection (**Figure 6E**). Unvaccinated mice lacking either T cells or B cells showed similar weight loss kinetics, with 80% T cell dep. mice and 20% B cell KO mice reaching humane endpoint at the same time as unvaccinated wildtype controls (**Figure 6E**). In comparison, i.n. vaccination with Tri:HuAd protected wildtype mice against SARS-CoV-2 (**Figure 6F**), further verifying our findings in **Figure 6B**. Of interest, vaccinated mice deficient in B cells showed no weight loss, similar to vaccinated wildtype controls (**Figure 6F**). In contrast, vaccinated mice lacking T cells experienced a transient moderate weight loss (10%), with 20% of them succumbing to infection (**Figure 6F**). On the other hand, i.n. vaccination with Tri:ChAd vaccine protected wildtype, T cell dep., and B cell KO animals equally well, as they did not show any weight loss throughout the course of experimentation (**Figure 6G**). These data indicate that the superior immunogenicity of i.n. Tri:ChAd vaccine is capable of compensating for the lack of either T cells or B cells.

Besides clinical outcomes, we further examined the role of B and T cells in control of viral burden in the lung 4-days post-infection (**Figure 6D**). Compared to wildtype hosts, lack of T or B cells in unvaccinated animals had little effects on lung viral burden (**Figure 6H, left**). Similar to the 2-day data (**Figure 6C**), i.n. Tri:HuAd vaccination moderately reduced viral loads in wildtype animals by approximately 2 log (**Figure 6H, middle**). However, lack of either T or B cells in Tri:HuAd vaccinated animals resulted in increased lung viral burden (∼1 log). Of interest, i.n. vaccination with Tri:ChAd vaccine led to complete clearance of virus at 4-days post-infection (**Figure 6H, right**). Similar to that observed with Tri:HuAd vaccine, lack of either T or B cells resulted in significantly increased lung viral titers (2-4 log), despite no changes in morbidity and mortality (**Figure 6G**).

Together, the above data indicate that first, single-dose intranasal immunization provides superior immune protection against SARS-CoV-2 over intramuscular immunization in a murine model with a mouse-adapted virus; secondly, Tri:ChAd vaccine is more potent than its Tri:HuAd counterpart; and thirdly, both humoral and T cell immunity contribute to protection provided by Ad-vectored respiratory mucosal immunization.

### Single-dose intranasal immunization with ChAd-vectored COVID-19 vaccine protects against lethal infection by SARS-CoV-2 variants of concern

Mouse-adapted strains of pandemic viruses such as SARS-CoV-2 MA-10 are indispensable tools for rapid assessment of prophylactic and therapeutic countermeasures, but by their very nature they are inherently dissimilar to bona fide SARS-CoV-2. Given the comprehensive protection observed with mouse-adapted virus, we next sought to assess the extent of vaccine-induced protection against wildtype SARS-CoV-2 including both ancestral and VOC strains of virus, which entailed the utilization of the K18-hACE2 mouse, a highly susceptible model extensively used for SARS-CoV-2 research that develops a rapid, lethal infection and cytokine storm traits observed in humans (Khoury et al., 2020). Since our data have collectively shown the superiority of the intranasal route of immunization with Tri:ChAd vaccine in both immunogenicity and protection, we focused on evaluating the protective potency of intranasal Tri:ChAd immunization in K18-hACE2 models of COVID-19.

Given that the bulk of our immunogenicity studies were carried out in BALB/c mice thus far, we began by assessing vaccine immunogenicity in K18-hACE2 mice (B6 background) and compared it to noncarrier wildtype C57BL/6 mice at 4-weeks following i.n. immunization with Tri:ChAd (**Figure S4A**). Similar to the neutralization titers observed in BALB/c hosts (**Figure 2D)**, sera from i.n. Tri:ChAd-immunized C57BL/6 animals showed robust SARS-CoV-2 neutralization (**Figure S4B**). Similarly, airway antigen-specific CD8^+^ T cell responses in C57BL/6 mice were also comparable (**Figure S4C**) to those in BALB/c counterparts (**Figure 3A/D/G**). Also, consistent with our findings in BALB/c mice (**Figure 5D/E**), there was markedly increased MHC II expression on airway macrophage populations (AM & IM) only following i.n. Tri:ChAd immunization (**Figure S4D**). Finally, upon comparing lung S-specific CD8 T cell responses, we saw CD8 T cell responses specific for ancestral SARS-CoV-2 spike antigens to be comparable to those for B.1.351 variant spike antigens (**Figure S4E**). These data suggest that intranasal Tri:ChAd vaccine is similarly immunogenic in genetically distinct mice as in BALB/c hosts and that T cell antigens are well conserved among ancestral and variant strains of SARS-CoV-2.

We then evaluated the protective efficacy of i.n. Tri:ChAd immunization in K18-hACE2 model following challenge with an ancestral strain of SARS-CoV-2. To this end, K18-hACE2 mice were immunized intranasally with a single-dose of Tri:ChAd vaccine. As controls, some mice were left unvaccinated (Naïve) or inoculated intranasally with an empty ChAd vector that does not express any SARS-CoV-2 antigens (ChAd:EV). Mice were then infected 4-weeks later with 1×10^5^ PFU of SARS-CoV-2/SB3 (**Figure 7A**), an early pandemic strain isolated in Toronto, Canada (Banerjee et al., 2020). Both naïve and ChAd:EV groups of animals quickly succumbed, reaching the clinical endpoint based on weight loss and/or neurological manifestations between 5-7 days post-infection (**Figure 7B/C**). In contrast, i.n. Tri:ChAd immunization largely prevented weight loss (**Figure 7B**) and mortality (**Figure 7C**), with 9/10 mice surviving infection. In agreement with the morbidity and mortality observations, while both naïve and ChAd:EV-treated animals had high lung viral burden, single-dose i.n. Tri:ChAd immunization provided sterilizing immunity within the lung (**Figure 7D**). These data indicate that a single-intranasal-dose of Tri:ChAd vaccine is sufficient to protect K18-hACE2 mice from lethal challenge with a wildtype ancestral human SARS-CoV-2 isolate.

**Figure 7.**
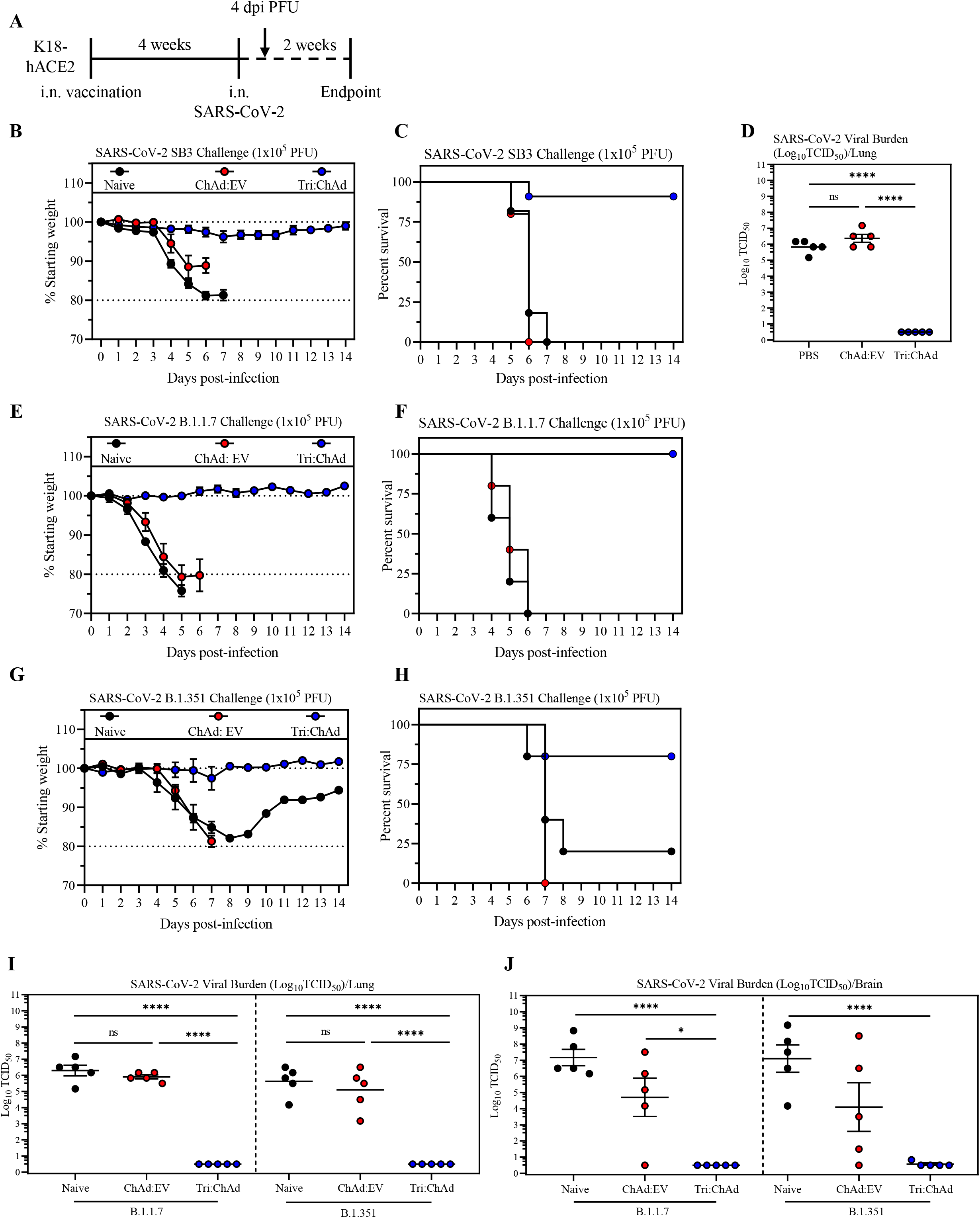
Intranasal Tri:ChAd immunization protects against lethal challenge with SARS-CoV-2 variants of concern. (A) Schema of SARS-CoV-2 challenge regimen. K18-hACE2 mice were intranasally (i.n.) vaccinated with a single dose of either EV:ChAd or Tri:ChAd for 4 weeks or left unvaccinated (naïve). Mice were infected i.n. with a lethal dose of ancestral SARS-CoV-2 SB3, variant B.1.1.7 or variant B.1.351. A subset of animals was sacrificed 4 days post-infection for enumeration of viral burden. (B) Changes in body weight over 2 weeks post-ancestral SARS-CoV-2 infection. (C) Survival of unvaccinated, i.n. EV:ChAd-treated or i.n. Tri:ChAd-vaccinated mice post-viral infection. (D) Viral burden (Log_10_TCID_50_) in the lung at 4 days post-infection. (E) Changes in body weight over 2 weeks post-SARS-CoV-2 B.1.1.7 infection. (F) Survival of unvaccinated, i.n. EV:ChAd-treated or i.n. Tri:ChAd-vaccinated mice post-B.1.1.7 infection. (G) Changes in body weight over 2 weeks post-SARS-CoV-2 B.1.351 infection. (H) Survival of unvaccinated, i.n. EV:ChAd-treated or i.n. Tri:ChAd-vaccinated mice post-B.1.351 infection. (I) Viral burden (Log_10_TCID_50_) in the lung at 4 days post-B.1.1.7 or B.1.351 infection. (J) Viral burden (Log_10_TCID_50_) in the brain at 4 days post-B.1.1.7 or B.1.351 infection. Data presented (B-J) represent mean ± SEM. Statistical analysis for panels D, I, and J were 1-way ANOVA with Tukey multiple comparisons test. Data is representative of 1-2 experiments, n=5-11 mice/group. *P < 0.05; ****P < 0.0001.

Different from earlier ancestral SARS-CoV-2 strains, the emerging variants of concern (VOC) have accumulated mutations in S, being able to evade the immunity elicited by first-generation vaccines (Harvey et al., 2021). This situation calls for developing next-generation vaccine strategies. Having observed superb protective efficacy of our next-generation i.n. Tri:ChAd vaccine strategy against lethal infection by both mouse-adapted (**Figure 6B/C/G/H**) and wildtype-ancestral (**Figure 7B/C/D**) strains of SARS-CoV-2, we evaluated its efficacy in K18-hACE2 model against two emerging VOC, B.1.1.7 and B.1.351. Compared to other VOC, B.1.351 is highly immune-evasive (Harvey et al., 2021) and indeed, while the immune serum from i.n. Tri:ChAd-vaccinated K18-hACE2 animals neutralized both ancestral and B.1.1.7 strains of virus equally well, it had considerably reduced capacity (MNT50: 70.70%) to neutralize B.1.351 variant (**Figure S4F**). Thus, K18-hACE2 mice were immunized intranasally with Tri:ChAd vaccine and control groups were set up and subsequently challenged with a lethal dose (1×10^5^ PFU) of the B.1.1.7 or B.1.351 variant as described in **Figure 7A**. Similar to what was observed with ancestral SARS-CoV-2 strain (**Figure 7B/C**), both unvaccinated naïve and ChAd:EV animals showed quick weight loss (**Figure 7E**), reaching the clinical endpoint between 5-6 days post-infection by B.1.1.7 variant (**Figure 7F**). In sharp contrast, single-dose i.n. immunization with Tri:ChAd vaccine completely protected the animals from morbidity and mortality (**Figure E/F**). Similarly, upon infection with B.1.351 variant, while 80% of unvaccinated naïve and all of ChAd:EV animals succumbed by 7-8 days, the vast majority of i.n. Tri:ChAd-immunized animals were well protected (**Figure 7G/H**). We next quantified viral burden in the lung at 4-days post-infection (**Figure 7A**). Since SARS-CoV-2 infection of K18-hACE2 mice also causes severe viremia and viral dissemination to the central nervous system (Zheng et al., 2021), we also assessed viral burden in the brain (**Figure 7J**). Unvaccinated naive mice had similarly high B.1.1.7 or B.1.351 viral burden within the lung (**Figure 7I**) and brain tissue (**Figure 7J**). Of interest, while ChAd:EV-inoculated mice had similarly high viral titers in the lung as in naïve mice (**Figure 7I**), they showed moderately reduced viral burden in the brain, compared to naïve mice (**Figure 7J**). Remarkably, mice intranasally immunized with Tri:ChAd developed sterilizing immunity against both B.1.1.7 and B.1.351 variants, showing no detectable viruses in either the lung (**Figure 7I**) or brain (**Figure 7J**). To further investigate the protective efficacy of i.n. Tri:ChAd vaccine against B.1.351 variant, we compared Tri:ChAd vaccine with a bivalent ChAd vaccine (Biv:ChAd) expressing only the N and RdRp antigens (**Figure S5A**). Following single-dose i.n. immunization and B.1.351 challenge, lung viral burden was quantified at 4-days (**Figure S5B**). Compared to unvaccinated controls, Biv:ChAd vaccine resulted in >25 times reduction in lung PFU whereas Tri:ChAd vaccine led to >75 times reduction (**Figure S5C**). Since Biv:ChAd vaccine does not express the S1 antigen, the data offers further support to the role of T cell immunity in neutralizing antibody-independent protection against SARS-CoV-2 (**Figure 6E/F/G/H**).

Collectively, the above data indicate that single-dose respiratory mucosal immunization with a multivalent next-generation ChAd-vectored COVID-19 vaccine induces robust, sterilizing immune protection against lethal infection by not only ancestral SARS-CoV-2 but importantly, highly virulent immune-evasive VOC. Furthermore, our data also suggest that Ad vector-induced trained innate immunity in the lung can broadly protect against systemic viral dissemination.

## Discussion

The effective global control of COVID-19 via immunization with first-generation vaccines has been threatened by the emerging VOC on top of the shortage of vaccine supplies that many countries are now facing (Harvey et al., 2021; Krause et al., 2021). This situation calls for the urgent development of not only next-generation vaccines but also diversified vaccine strategies. In response, we have developed adenoviral (Ad)-vectored trivalent COVID-19 vaccines for respiratory mucosal route of delivery. Our next-generation vaccine strategy is designed to be effective against both ancestral and variants of SARS-CoV-2 via its expression of the original S1 antigen and the highly conserved T cell antigens N and RdRp, and induction of all-around protective mucosal immunity consisting of antibody and T cell immunity as well as trained innate immunity (Jeyanathan et al., 2020). Indeed, by using murine models our current study has provided strong evidence that the respiratory mucosal route of immunization is superior to intramuscular immunization at inducing neutralizing antibodies, mucosal tissue-resident memory T cells and trained airway resident macrophages. We further show that the choice of Ad vector also is of importance, with chimpanzee-derived Ad68 platform (Tri:ChAd) outperforming the human Ad5 counterpart (Tri:HuAd). We have also provided the evidence that both B and T cell immunity is required for optimal protection and that trained innate immunity is involved in limiting systemic viral dissemination. Thus, a single intranasal, but not intramuscular, dose of Tri:ChAd vaccine provides robust protective immunity against infection by a mouse-adapted SARS-CoV-2, an ancestral pandemic strain or two VOC including the highly immune-evasive B.1.351 in wildtype BALB/c and K18-hACE2 mouse models.

To the best of our knowledge, our study represents the first to demonstrate the *in vivo* protective efficacy of a novel, multivalent next-generation vaccine strategy against both ancestral SARS-CoV-2 and emerging VOC in animal models. Although a recent study shows the ability of a first-generation S-encoding mRNA vaccine (CVnCoV) to protect from B.1.351 variant infection in murine models (Hoffmann et al., 2021a), this candidate vaccine has recently reported disappointing efficacy results from clinical trials. It is widely accepted that the next-generation vaccine strategies ought to take into consideration both vaccine multivalency and route of delivery (Jeyanathan et al., 2020; Teijaro and Farber, 2021). Almost all first-generation recombinant COVID-19 vaccines were designed for i.m. administration and to express only the S protein. As the COVID-19 vaccine effort intensifies, there has been a growing interest in investigating the protective immunity by respiratory mucosal delivery of first-generation viral-vectored vaccines in preclinical models (Bricker et al., 2021; Cao et al., 2021; Hassan et al., 2020, 2021a; Ku et al., 2021). However, the majority of these studies did not compare the intranasal route of immunization with the intramuscular route, nor did they test the protection conferred by first-generation vaccine candidates against the emerging VOC. The studies reported by a group of scientists at WUSTL are the only ones where the intranasal route was compared with intramuscular immunization (Bricker et al., 2021; Hassan et al., 2020) and this group also extended their study to demonstrate the ability of intranasal ChAd-S immunization to protect against a Washington strain of virus displaying B.1.351 spike protein (Hassan et al., 2021b). By comparison, with the goal of developing next-generation COVID-19 vaccine strategies, we have bioengineered two different Ad-vectored trivalent vaccines (Tri:HuAd & Tri:ChAd) and extensively compared their immunogenicity and protective efficacy against ancestral and emerging variant strains of SARS-CoV-2 following single-dose intramuscular or intranasal route of immunization. Our preclinical findings support the single-dose respiratory mucosal delivery of a trivalent ChAd-vectored vaccine to be the most effective next-generation COVID-19 vaccine strategy. Our study thus provides the important proof of concept for its further clinical development. If proven successful, the next-generation vaccine strategies such as ours may be deployed as a booster to bolster mucosal protective immunity against emerging VOC and to extend the durability of protective immunity following immunization with first-generation vaccines (Jeyanathan et al., 2020).

The superiority of intranasal COVID-19 immunization at inducing both protective humoral and T cell mucosal immunity over the intramuscular route observed in our current study is well aligned with the established paradigm associated with other vaccine strategies (Belyakov and Ahlers, 2009; Jeyanathan et al., 2018; Neutra and Kozlowski, 2006). It has also been observed in murine and hamster models of COVID-19 using a ChAd-vectored first-generation vaccine (Bricker et al., 2021; Hassan et al., 2020). The high degree of tissue compartmentalization of immunity dictated by the route of immunization is not only limited to animal models. It was observed that inhaled aerosol MVA TB vaccine induced respiratory mucosal T cell responses in humans whereas intradermal injection of the same viral-vectored vaccine failed to do so (Satti et al., 2014). The findings from our recently completed phase 1b trial study comparing inhaled aerosol HuAd-vectored TB vaccine with intramuscular delivery lend further support to this principle. Given that all of the currently approved viral-vectored COVID-19 vaccines including ChAdOx1nCov-19 (AstraZeneca/Oxford), Ad26.COV2-S (J&J), Gam-COVID-Vac (Gamaleya) and Ad5-nCoV (CanSino) are intramuscularly administered, they are expected to be unable to induce protective respiratory mucosal immune mechanisms including tissue-resident memory T cells and tissue-resident trained innate immunity (Jeyanathan et al., 2020). We have also recently shown that SARS-CoV-2-specific IgA, which is enriched at mucosal surfaces like the lung and is potently induced by mucosal vaccine, can induce neutrophils to undergo NETosis. These NETs are capable of trapping and killing virus, thereby limiting spread (Stacey et al., 2021). These limitations of i.m. delivery, along with our current findings and those from others (Bricker et al., 2021; Hassan et al., 2020), should further bolster the global effort in developing respiratory mucosal-deliverable next-generation COVID-19 vaccines. In this regard, there have been at least two clinical trials testing either inhaled aerosol ChAdOx1nCov-19 (Singh et al., 2021) or intranasally delivered ChAd-SARS-CoV-2-S developed by scientists at WUSTL (Clinical Trial NCT04751682).

Our current study has also provided the novel experimental evidence that both B and T cells are required for optimal protection. Of note, our data suggest that the clinical outcomes (body weight losses and mortality) may not always corroborate with viral burden, depending on both the immune pathway and relative potency of vaccine. We have observed that while Tri:HuAd-vaccinated B cell-deficient hosts well-protected in terms of clinical outcomes suffered impaired lung viral clearance, T cell-deficient hosts suffered both moderately worsened clinical outcomes and lung viral clearance. On the other hand, the robust i.n. Tri:ChAd vaccine strategy protected wildtype, B cell- and T cell-deficient hosts equally well in terms of clinical outcomes, whereas lack of B or T cells led to impaired lung viral clearance. Besides using T cell-depletion approach, the role of vaccine-induced T cell immunity in protection is supported further by our observations from B.1.135-infected K18-hACE2 animals immunized with a bivalent, S antigen-independent N-RdRp-expressing ChAd vaccine. These findings together further support the relevance of broadening the breadth of T cell immunity in COVID-19 vaccine design.

When recombinant Ad-vectored vaccines are delivered via the respiratory mucosal route to humans, it helps bypass the pre-existing anti-vector immunity which is much more prevalent in the circulation than in the respiratory tract. This is particularly relevant to the use of human Ad5 and Ad26 vectors (Jeyanathan et al., 2020). Nonetheless, ChAd-vectored vaccines have an advantage over both Ad5 and Ad26 vectors in that humans have little pre-existing immunity against ChAd viruses. Experimentally we have previously shown that intranasal ChAd-vectored vaccine was not impacted by high levels of anti-Ad5 immunity in murine lungs (Jeyanathan et al., 2015). Besides these potential advantages in respiratory mucosal application of ChAd-vectored COVID-19 vaccine in humans, our current study provides strong evidence that ChAd-vectored trivalent COVID-19 vaccine is also much more immunogenic and protective than HuAd-vectored counterpart, even when a smaller dose was given.

In summary, in response to the pressing need to develop next-generation vaccine strategies against the emerging VOC of SARS-CoV-2, we have developed and extensively evaluated two Ad-vectored vaccines for their immunogenicity and protective efficacy in relevant murine models established with mouse-adapted, wildtype ancestral, and emerging variant strains of virus. Our next-generation vaccine strategy is unique in both vaccine design and route of delivery. Our findings demonstrate that firstly, respiratory mucosal route is superior to intramuscular route of immunization with trivalent Ad-vectored COVID-19 vaccine; secondly, ChAd-vectored vaccine is more potent than HuAd-vectored counterpart; thirdly, optimal protective immunity requires both B and T cells; and lastly, Ad vector-induced trained innate immunity in the lung helps limit systemic viral dissemination. Thus, a single-dose intranasal immunization with trivalent ChAd-vectored vaccine induced robust protection against lethal infection by not only ancestral, but also emerging variant strains of SARS-CoV-2. These exciting findings warrant further clinical studies to evaluate inhaled aerosol delivery of our multivalent viral-vectored COVID-19 vaccine to the respiratory tract in humans.

## Acknowledgments

The work was supported by the Canadian Institutes of Health Research (CIHR) COVID-19 Rapid Research Project (ZX et al), CIHR Foundation Program (ZX), the Innovative Research Program of National Sanitarium Association of Canada (ZX), CIHR New Investigator Award and Ontario Early Research Award (MSM), CIHR Canada Graduate Scholarship Doctoral Award and Physicians’ Services Incorporated Research Trainee Fellowship (AZ), Ontario Graduate Scholarship (MRD), Ontario Graduate Scholarship and Canadian Society for Virology United Supermarket Studentship (HDS), Canadian Foundation for Innovation, Ontario Government, and McMaster University. Authors are thankful to Turnstone Biologics Inc for provision of ChAd68 vector, to Dr. Ralph Baric for provision of a mouse-adapted SARS-CoV-2 (MA-10) and to Dr. Carolina Ilkow for provision of subcloning materials.

## Author contributions

SA, MRD, YW, KM, MJ, FS, BDL, MSM, and ZX conceived and designed the study. SA, MRD, AZ, HDS, AM, AK, RS, JB, GY, XL, FW, JCA, AZ, and AG performed experiments. JFEK, AP, and MJ designed and constructed RBD tetramers. SA, MRD, AZ, HDS, AM, FW, and JCA analyzed data. US, NK, MFM, SA, and BDL designed, constructed and purified vaccines. SA, MRD, MSM, and ZX wrote the paper.

## Supplemental Figure Titles and Legends

**Figure S1.**
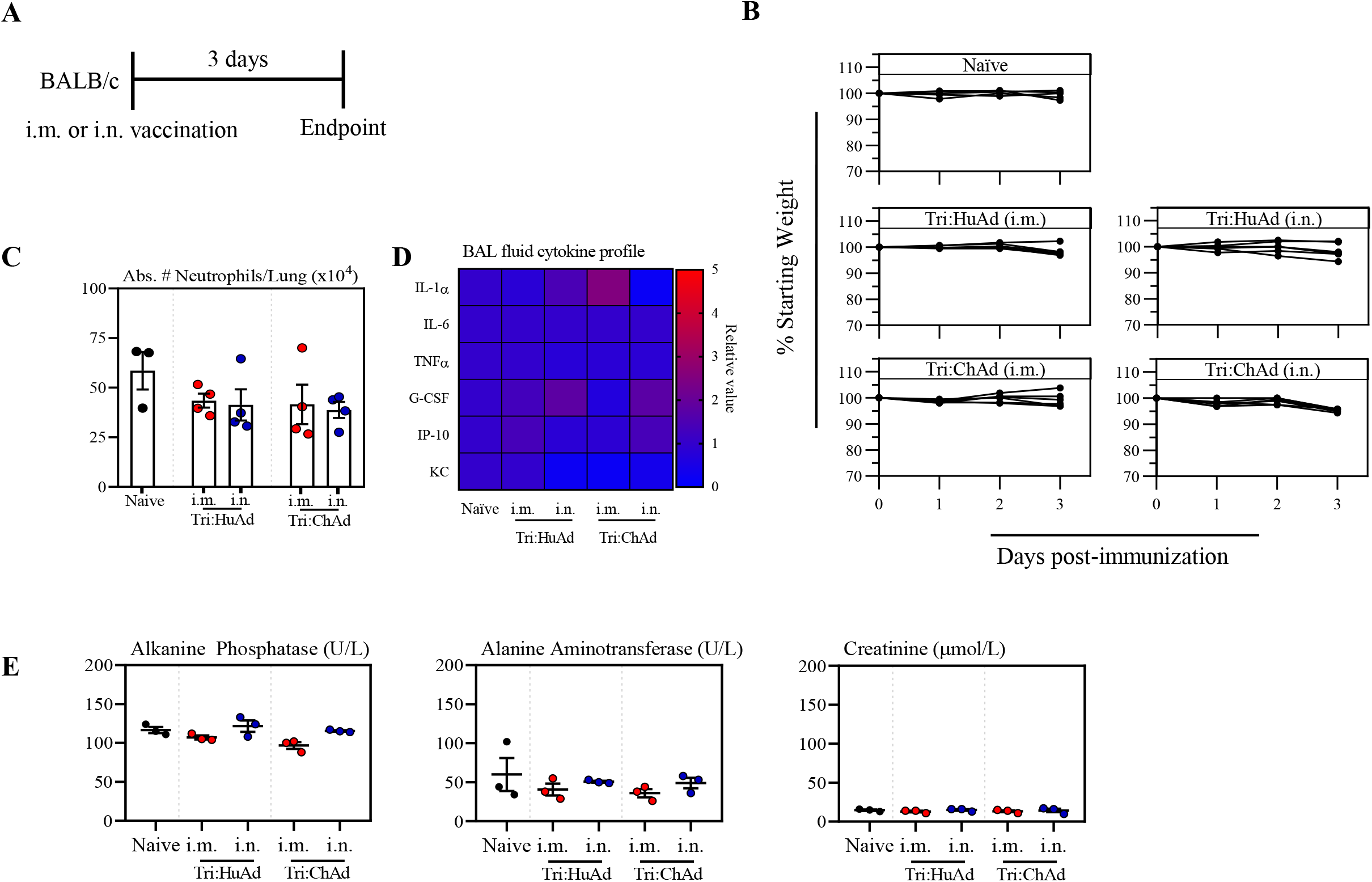
Acute safety assessment of intramuscularly or intranasally administered Tri:HuAd and Tri:ChAd COVID-19 vaccines. (A) Schema of vaccination regimen. BALB/c mice were either intramuscularly (i.m.) or intranasally (i.n.) vaccinated with a single dose of either Tri:HuAd or Tri:ChAd. Animals were sacrificed 3 days post-immunization for inflammatory assessment. (B) Changes in body weight over 3 days post-vaccination. (C) Absolute number of neutrophils in the lung at 3 days post-immunization. (D) Cytokine levels in bronchoalveolar lavage fluids at 3 days post-immunization. (E) Serum levels of biomarkers for hepatotoxicity and nephrotoxicity at 3 days post-immunization. Data presented represent mean ± SEM from n=3 mice/group.

**Figure S2.**
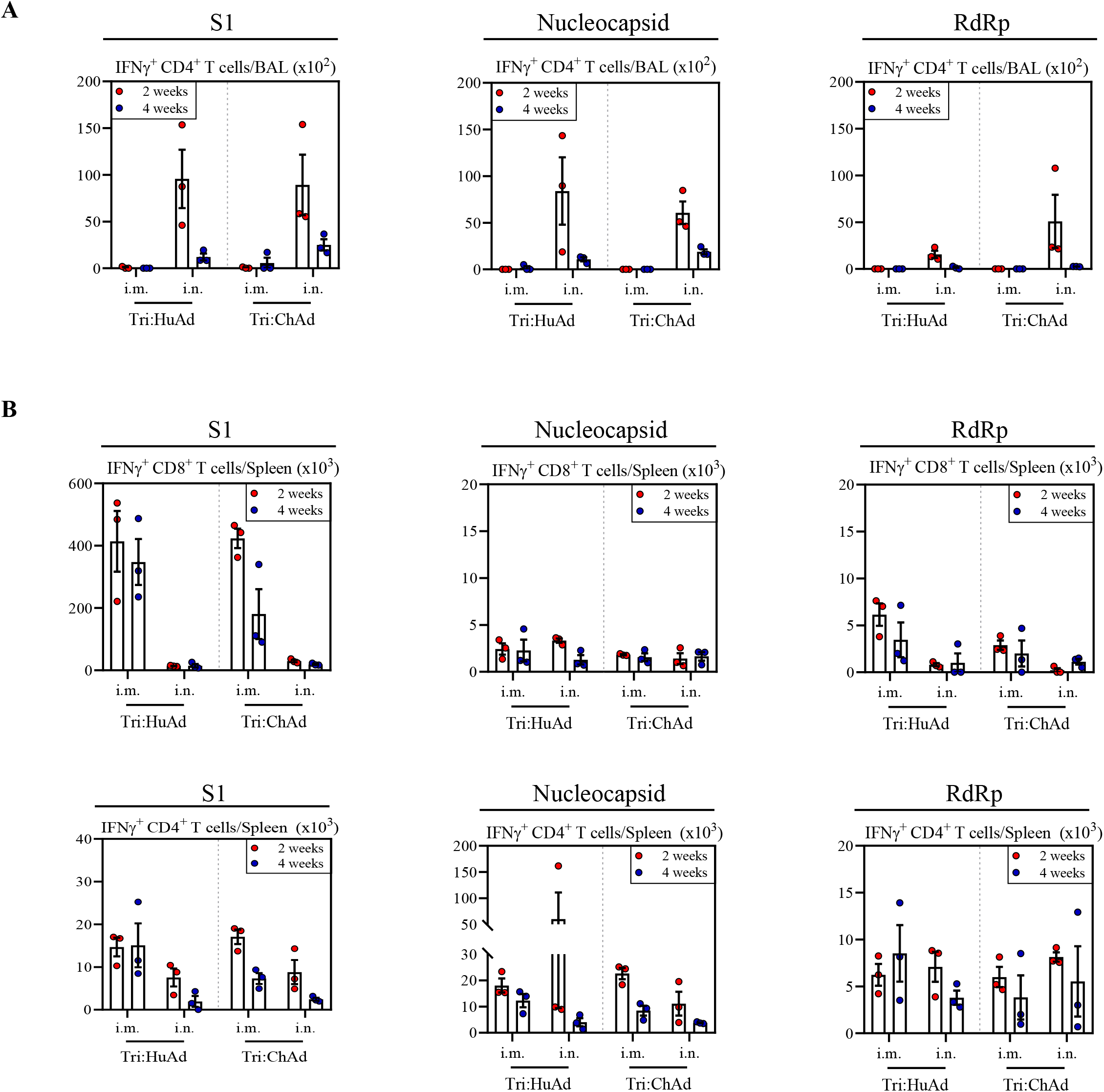
Comparison of antigen-specific CD4 and CD8 T cells in BAL and spleen following single-dose immunization with Tri:HuAd or Tri;ChAd vaccine. (A) Absolute numbers of antigen-specific IFNγ^+^CD4^+^ T cells in the airway at 2 (red) and 4 (blue) weeks post-immunization. (B) Absolute numbers of antigen-specific IFNγ^+^CD8^+^ T cells (top panels) and antigen-specific IFNγ^+^CD4^+^ T cells (bottom panels) in the spleen at 2 (red) and 4 (blue) weeks post-immunization. Data presented represent mean ± SEM from n=3 mice/group.

**Figure S3.**
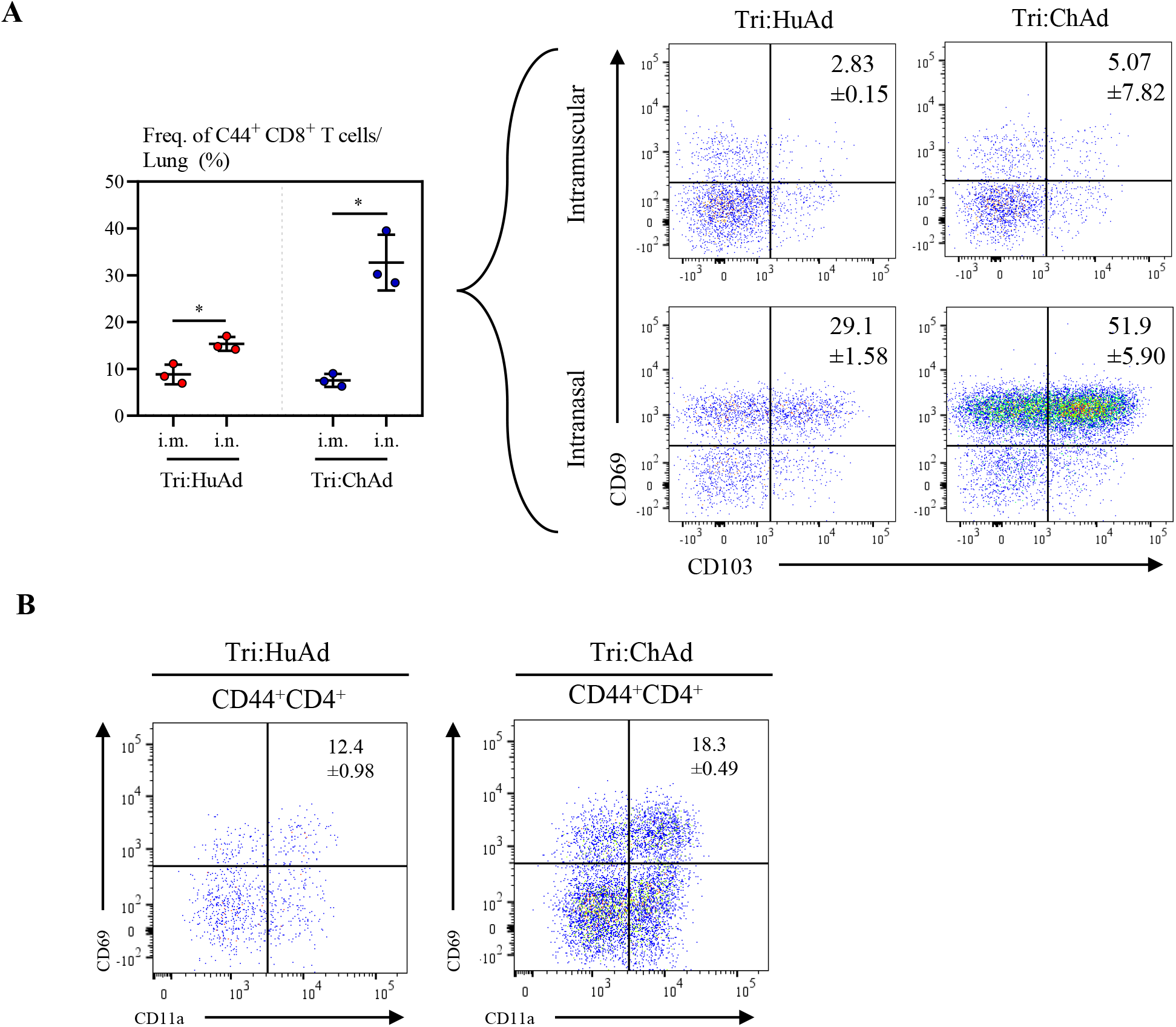
Comparison of lung tissue-resident memory CD8 and CD4 T cells following single-dose immunization with Tri:HuAd or Tri:ChAd vaccine. (A) Frequency of CD44^+^CD8^+^T cells in the lung (left panel), dot plots showing frequencies of T_RM_ markers on CD44^+^CD8^+^ T cells in the lung (right panels) at 8 weeks post-immunization. (B) Dot plots showing frequencies of T_RM_ markers on CD44^+^CD4^+^ T cells in the lung at 8 weeks post-immunization. Data presented in A represent mean ± SEM from n=3 mice/group. Statistical analysis for panel A was 1-way ANOVA with Tukey multiple comparisons test. *P < 0.05.

**Figure S4.**
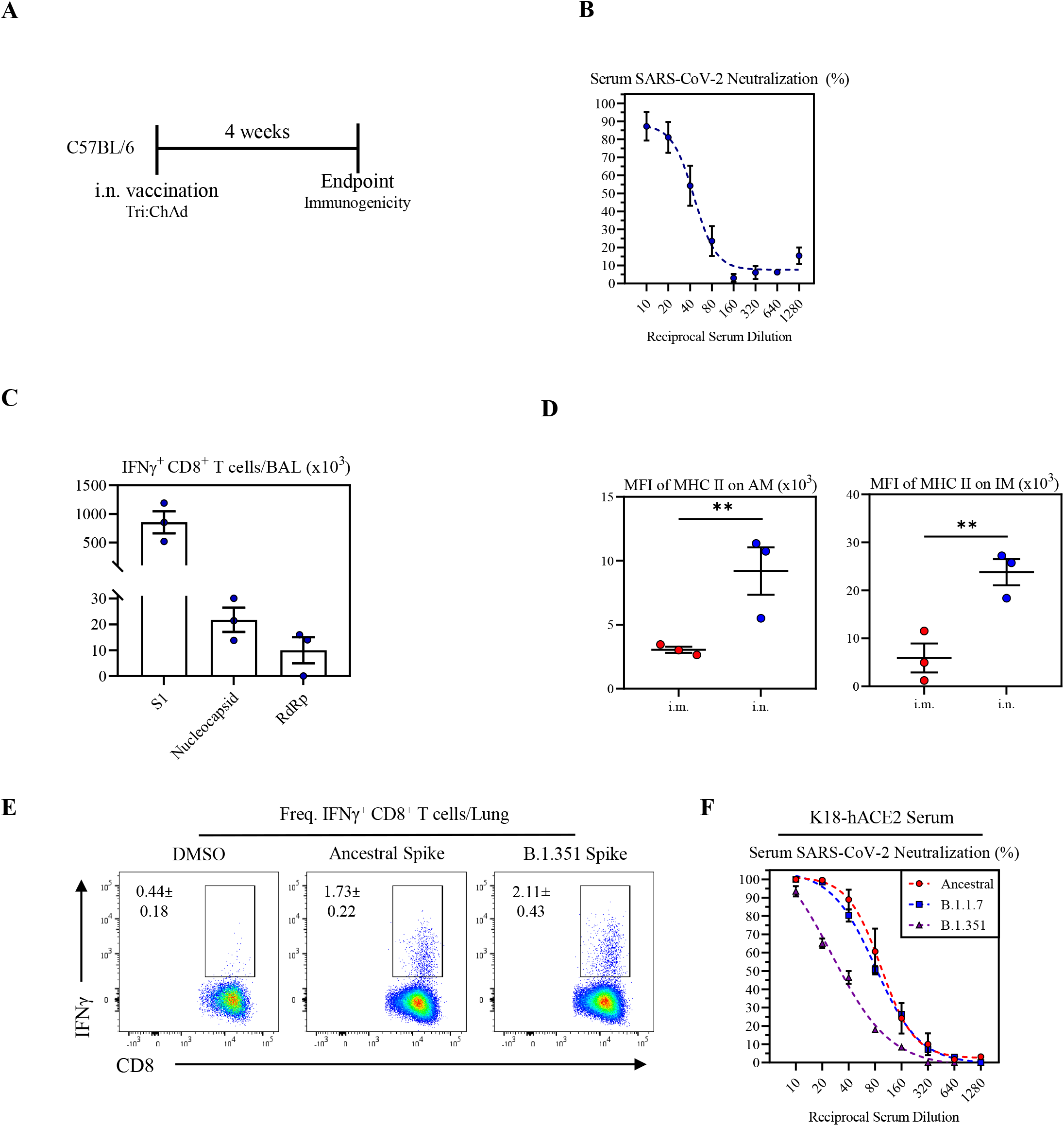
Characterization of immunogenicity of intranasal immunization with Tri:ChAd vaccine in wildtype C57BL6 or K18-ACE2 mice. (A) Schema of vaccination regimen. C57BL/6 or K18-ACE2 mice were intranasally (i.n.) vaccinated with a single dose of Tri:ChAd for 4 weeks. (B) Serum neutralizing antibody responses at 4 weeks post-immunization of C57BL/6 mice, measured by percent (%) neutralization utilizing a live SARS-CoV-2 microneutralization (MNT) assay. (C) Absolute number of antigen-specific IFNγ^+^ CD8^+^ T cells in the airway at 4 weeks after i.n. Tri:ChAd immunization in C57BL/6 mice. (D) MFI of MHC II expression on AM (left panel) and IM (right panel) in BAL at 4 weeks post-immunization. (E) Dot plots showing frequencies of spike-specific IFN γ^+^ CD8^+^ T cells in lung mononuclear cells upon stimulation with either ancestral or variant SARS-CoV-2 spike protein at 4 weeks post-immunization. (F) K18-hACE2 mice were vaccinated i.n. with Tri:ChAd for 4 weeks and sera were collected. Serum neutralizing antibody responses measured by percent (%) neutralization utilizing a live SARS-CoV-2 ancestral (red), B.1.1.7 (blue), or B.1.351 (purple) microneutralization (MNT) assay. Data presented in B,C,D,E,F represent mean ± SEM from 3-5 mice/group. Statistical analysis for panel D was 1-way ANOVA with Tukey multiple comparisons test. **P < 0.01.

**Figure S5.**
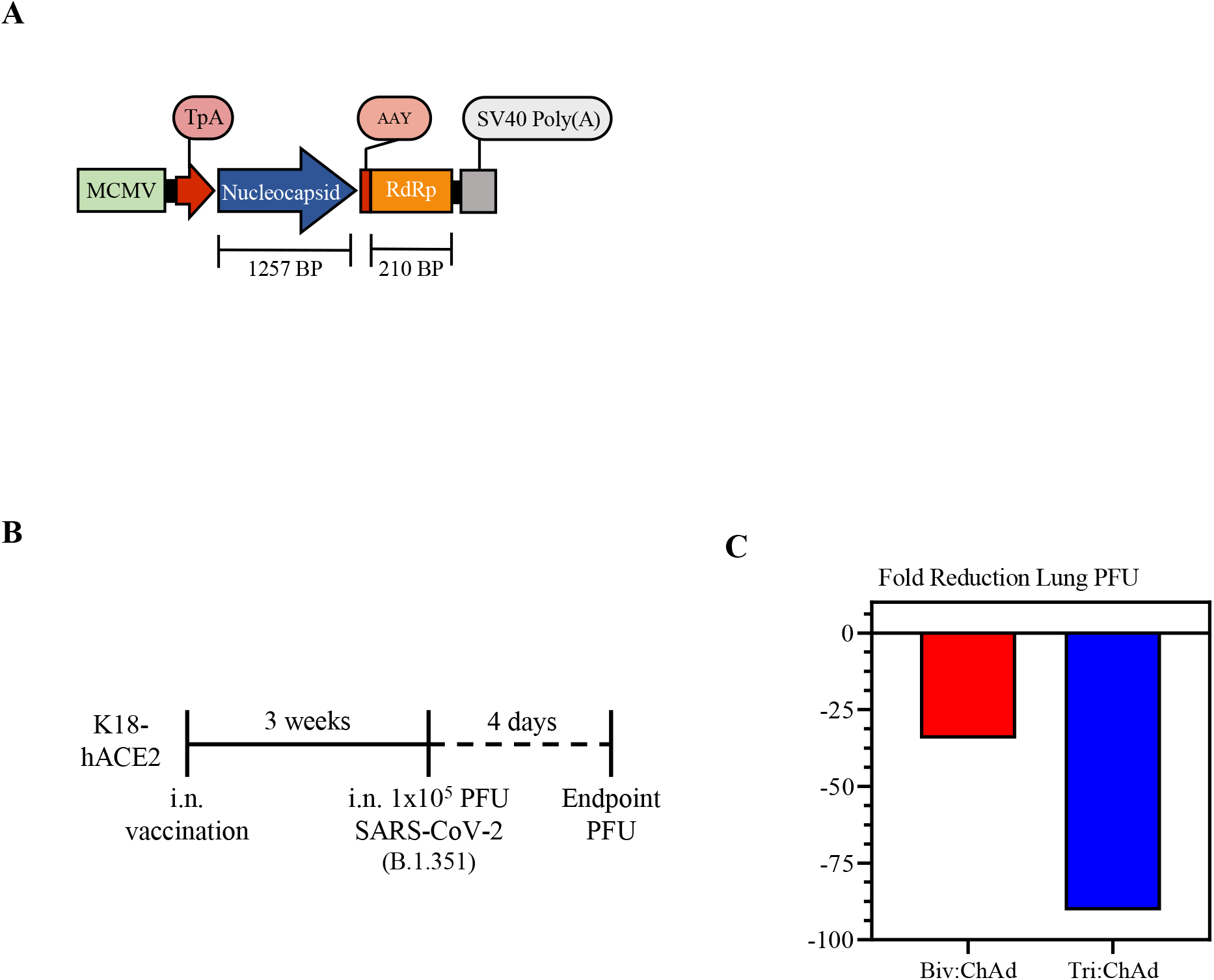
Construction and protective efficacy of ChAd-vectored bivalent COVID-19 vaccine. A. Transgene cassette diagram. The transgene construct is under control of the MCMV promoter. Nucleocapsid and RdRp are expressed as a polyprotein. Construction of Biv:ChAd is identical to Tri:ChAd but lacks S1-VSVG. B. Schema of SARS-CoV-2 challenge regimen. K18-hACE2 mice were intranasally (i.n.) vaccinated with a single dose of either Biv:ChAd or Tri:ChAd for 3 weeks or left unvaccinated. Mice were infected i.n. with a lethal dose of SARS-CoV-2 variant B.1.351. Animals were sacrificed 4 days post-infection for assessment of lung viral burden. C. Fold reduction in lung PFU in Biv:ChAd (red) or Tri:ChAd (blue) immunized animals, relative to unvaccinated controls. Data in C represent the average value from 3 mice/group.

## STAR METHODS

### CONTACT FOR REAGENT AND RESOURCE SHARING

Further information and requests for resources and reagents should be directed to and will be fulfilled by the Lead Contacts, Dr. Zhou Xing and Dr. Matthew Miller.

### EXPERIMENTAL MODEL AND SUBJECT DETAILS

#### Mice

Age-matched 6–8-wk-old wild-type female BALB/c, C57BL/6J, or B6.Cg-Tg (K18-ACE2) 2Prlmn/J mice were purchased from either Charles River Laboratories (Saint Constant, QC, Canada) or The Jackson Laboratory (Bar Harbor, ME, United States). B cell-deficient mice C.Cg-*Igh-J^tm1Dhu^* of Balb/c background were purchased from Taconic Biosciences (Germantown, NY, United States). Animals were housed in either a specific pathogen-free level B or a Containment Level 3 Facility at McMaster University, Hamilton, ON, Canada. All experiments were performed in accordance with institutional guidelines from the Animal Research and Ethics Board.

### METHOD DETAILS

#### Vaccine construction

The transgene cassette was constructed through a series of overlapping polymerase chain reactions (PCR) wherein transgene expression is under control of the murine CMV (mCMV) promoter and protein translation is initiated with the human tissue plasminogen (tPA) signal sequence (**Figure 1A**). The first overlapping PCR product contained the tPA signal sequence upstream of the S1 sequence of the Wuhan-Hu-1 Isolate of SARS-CoV-2 (GenBank: MN908947.3) fused to the vesicular stomatitis virus G protein transmembrane (VSVG TM) domain to facilitate trimerization and exosome targeting. This PCR product was cloned in pCY1 plasmid which contains the mCMV promoter. The second overlapping PCR product contained the porcine teschovirus-1 2A (P2A) skip sequence upstream of the full-length nucleocapsid sequence from the same SARS-CoV-2 isolate fused to a highly conserved region of nsp12 (RNA-dependent RNA polymerase (RdRp)).

The sequence of RdRP was chosen based on conserved sequence homology to bat coronaviruses and further refined to include several predicted high affinity human CD8 T cell epitopes on HLA 0101, 0201, and 0301. The second overlapping PCR product was cloned downstream of the VSVG TM domain in pCY1 to generate the complete expression cassette. The same transgene cassette was cloned in the shuttle plasmids used during co-transfection to rescue the tri-valent, replication-defective human serotype 5 adenoviral-vectored (Tri:HuAd) and chimpanzee serotype 68 adenoviral-vectored (Tri:ChAd) COVID-19 vaccines.

Tri:HuAd was packaged and rescued in HEK293 cells through a two-plasmid co-transfection system as previously described (Wang et al., 2004). Tri:ChAd was also constructed and rescued in HEK293 cells via direct subcloning or similarly through a two-plasmid co-transfection system. Briefly, the transgene cassette was PCR amplified to incorporate restriction enzyme sites and cloned in a shuttle vector containing a unique FspI cut site. The shuttle vector was then linearized with FspI and used for co-transfection with an SrfI-linearized plasmid containing the E1/E3-deficient ChAd68 genomic backbone. Both trivalent vaccines were further amplified in HEK293 cells and subsequently purified by cesium chloride density gradient ultracentrifugation.

#### Cell lines and SARS-CoV-2 viruses

Vero E6 (CRL-1586, American Type Culture Collection (ATCC), Manassas, VA, United States) were cultured at 37°C in Dulbecco’s Modified Eagle medium (DMEM) supplemented with 10 % fetal bovine serum (FBS), 1 % HEPES pH7.3, 1 mM sodium pyruvate, 1% L-Glutamine and 100 U/mL of penicillin–streptomycin. SARS-CoV-2 strain SB3-TYAGNC was provided by Dr. Arinjay Banerjee, Dr. Karen Mossman, Dr. Samira Mubareka, and Dr. Rob Kozak and isolated as described previously (Banerjee et al., 2020). SARS-CoV-2 strain MA-10 was generously provided by Dr. Ralph Baric (Leist et al., 2020). SARS-CoV-2 strain hCoV-19/England/204820464/2020 (B.1.1.7 variant, NR-54000, Public Health England) and strain hCoV-19/South Africa/KRISP-K005325/2020 (B.1.351 variant, NR-54009, African Health Research Institute) were both obtained from BEI Resources (Manassas, VA, United States).

#### Immunization and infection

Animals were anesthetized with isoflurane and vaccinated with 5×10^7^ PFU of a recombinant human adenovirus (Ad) serotype 5 SARS-CoV-2 vaccine (Tri:HuAd) or 1×10^7^ PFU of a recombinant chimpanzee adenovirus serotype 68 SARS-CoV-2 vaccine (Tri:ChAd). Where indicated, an empty Ad vector (the same adenoviral backbone lacking the vaccine transgene) was included as a control. Intranasal vaccinations were performed with a final volume of 25µL diluted in PBS. Intramuscular vaccinations were performed with a final volume of 100µL administered in equal volumes into the quadricep muscle of each hind leg. Infections were carried out with 1×10^5^ PFU SARS-CoV-2 administered intranasally in a final volume of 40µL diluted in PBS. Mice were monitored for clinical signs and weight loss daily, with 80% of initial weight considered humane endpoint, in accordance with institutional guidelines.

#### *In vivo* T cell depletion

T cell depletion was carried out utilizing previously published and validated protocols (Yao et al., 2018). 200 µg of Anti-CD4 (clone GK1.5) and anti-CD8 (clone 2.43) depleting, or an IgG isotype control antibodies (MilliporeSigma, Etobicoke, ON, Canada) were intraperitoneally administered as a single bolus 26-days post-vaccination. A second 100 µg dose was administered 2-days following the first dose, and repeated every 8-days to maintain depletion, as per experimental requirements.

#### Bronchoalveolar lavage, lung, mediastinal lymph node, nasal turbinate and spleen mononuclear cell isolation

Mice were euthanized by exsanguination. Cells from bronchoalveolar lavage (BAL), lung tissue, spleen, and mediastinal lymph nodes (MLN) were isolated as previously described (D’Agostino et al., 2020; Jeyanathan et al., 2015; Yao et al., 2018). Briefly, BAL was performed by instillation with 250 µL, followed by 200 µL of PBS. This fraction was utilized for downstream soluble factor analysis. Further instillation of 3x 300 µL of PBS was performed for BAL cell retrieval. Lungs were minced into small pieces and digested with collagenase type 1 (ThermoFisher Scientific Waltham, MA, United States) at 37°C in an agitating incubator. A single cell suspension was obtained by crushing the digested tissue through 100 µm basket filter (BD Biosciences, San Jose, CA, United States), with red blood cells being lysed with Ammonium-Chloride-Potassium (ACK) buffer for 2 minutes. Nasal turbinates, MLN, and spleen were ground between frosted glass microscope slides prior to passing through a 100 µm basket filter. Isolated cells were resuspended in complete RPMI 1640 (10 % FBS, 1% L-glutamine, 100 U/mL penicillin/streptomycin, 1% HEPES pH 7.3, 1% MEM non-essential amino acids (Gibco, Gaithersburg, MD, United States), 1% sodium-pyruvate (Gibco, Gaithersburg, MD, United States). Cell numbers were quantified in Turk’s Blood Dilution Fluid (RICCA Chemical, Arlington, TX, United States) and counted under a microscope. Where required, cells were counted automatically by a Sceptre 3.0 Cell Counter and Software Pro (Millipore Sigma, Etobicoke, ON, Canada).

#### Transgene protein analysis by Western blot

A549 cells (CCL-185, ATCC, Manassas, VA, United States) were cultured at 37°C in DMEM supplemented with 10% FBS, 1 % HEPES pH 7.3, 1% L-glutamine and 100 U/mL of penicillin–streptomycin. Cells were seeded in 24-well plates (7.5×10^4^ cells/well) 24 hours prior to infection. Infections were carried out with Tri:HuAd (MOI 100) and Tri:ChAd (MOI 50) diluted in PBS (with Mg^2+^ and Ca^2+^). 18 hours post-infection, cells were lysed using RIPA buffer (VWR, Mississauga, ON, Canada) containing a protease inhibitor cocktail (ThermoFisher Scientific Waltham, MA, United States). Lysates were quantified using a BCA kit (ThermoFisher Scientific Waltham, MA, United States) and 60 µg of each lysate was boiled at 98°C with 1X sample buffer (6.35% v/v 1M Tris, pH 6.8, 46.5% v/v 10X SDS, 20% v/v glycerol, and 5% v/v β-mercaptoethanol; MilliporeSigma, Etobicoke, ON, Canada) for 10 minutes. The samples were run on a 4-12% SDS-PAGE gel (ThermoFisher Scientific Waltham, MA, United States) for 1.5 hours at 100 V and transferred to nitrocellulose membrane (VWR, Mississauga, ON, Canada) using wet-transfer at 125 mA for 1.5 hours. The membrane was incubated with primary antibodies (1:1000 Anti-VSV-G (ThermoFisher Scientific Waltham, MA, United States), 1:2000 Anti-NP (ThermoFisher Scientific Waltham, MA, United States) and 1:5000 GAPDH (MilliporeSigma, Etobicoke, ON, Canada) diluted in 5% skim milk, followed by anti-mouse and anti-rabbit IRDye secondary antibodies (LI-COR, Lincoln, NE, United States) diluted in 5 % skim milk. The membrane was developed using an Odyssey CLx (LI-COR, Lincoln, NE, United States).

#### Recombinant antigen production

Plasmids encoding mammalian cell codon optimized sequences for the receptor binding domain (RBD) and full-length spike of SARS-CoV-2 was generously gifted from the lab of Dr. Florian Krammer (Amanat et al., 2020)(Icahn School of Medicine, NY, United States). Proteins were produced in Expi293F cells (ThermoFisher Scientific Waltham, MA, United States) according to the manufacturers’ instructions and purified as previously described (Stadlbauer et al., 2021). Briefly, when culture viability reached 40%, supernatants were collected and spun at 500 x g for 5 minutes. The supernatant was then incubated by shaking overnight at 4°C with 1 mL of Ni-NTA agarose (Qiagen, Germantown MD, United States) per 25 mL of transfected cell supernatant. The following day 10 ml polypropylene gravity flow columns (Qiagen, Germantown, MD, United States) were used to elute the protein. Recombinant RBD was concentrated in a 10 kDa Amicon centrifugal units (Millipore Sigma, Etobicoke, ON, Canada), and recombinant Spike was concentrated in a 50kDa Amicon centrifugal unit (Millipore Sigma, Etobicoke, ON, Canada) prior to being resuspended in phosphate buffered saline (PBS).

#### RBD tetramer construction

The recombinant RBD B cell tetramer was produced through biotinylation and tetramerization as previously described (Taylor et al., 2012). A decoy tetramer was created as previously described (Taylor et al., 2012), to gate out non-RBD binding B cells. The decoy tetramer was constructed through conjugating streptavidin-PE to Alexa Fluor 647 (Thermo Fisher Scientific) for 1 hour at room temperature. Excess Alexa Fluor 647 was removed through washing and centrifugation with 100 kDa Amicon spin columns (MilliporeSigma, Etobicoke, ON, Canada). The solution was then incubated with an irrelevant biotinylated protein at sixfold molar excess for 30 minutes at room temperature. The concentration of the resulting decoy tetramer was calculated by the absorbance of PE at 565 nM and diluted to 1 µM.

#### Enzyme Linked Immunosorbent Assay (ELISAs) for antibody measurement

96-well NUNC-MaxiSorp^TM^ plates (Thermo Scientific, Waltham, MA, United States) were coated overnight at 4°C with SARS-CoV-2 RBD, or full-length spike, diluted to 2 µg/mL in bicarbonate-carbonate coating buffer (pH 9.4). Plates were blocked by shaking for 1 hour at 37°C with reagent diluent (0.5% bovine serum albumin (BSA), 0.02 % sodium azide, in 1X Tris-Tween buffer).

Samples were serially diluted from 1:10 (serum), 1:4 (BAL), or a 1:2 (nasal wash, nasal turbinate, cecum) starting dilution. BAL, nasal wash, and nasal turbinate samples were first concentrated through Pierce^TM^ Protein Concentrators with a 50 kDa molecular weight cut-off (MWCO) (ThermoFisher Scientific Waltham, MA, United States) according to the manufacturer’s instructions, with volumes normalized prior to concentration. Samples were arranged such that one row contained only antigen and secondary antibodies and served as the plate blank. Following a 1 hour incubation with shaking at 37°C, plates were washed three times with 1X Tris-Tween wash buffer. After washing, goat anti-mouse-biotin antibodies (Southern Biotech, Birmingham, AL, United States) IgA (1:2000), IgG (1:5000), IgG1 (1:5000), IgG2a (1:5000)) were diluted in reagent diluent and added to all wells. Plates were again incubated for 1 hour, with shaking, at 37 °C, followed by three washes with 1X Tris-Tween buffer. A streptavidin-alkaline phosphatase secondary antibody (1:2000, Southern Biotech, Birmingham, AL, United States) was added to all wells for 1 hr with shaking at 37°C. Plates were subsequently washed three times prior to addition of pNPP one component microwell substrate solution (Southern Biotech, Birmingham, AL, United States) to each well. Plates were developed for 10 minutes and the reaction was quenched with an equal volume 3N sodium hydroxide. The optical density (O.D.) at 410 nm was read on a SpectramaxI3 (Molecular Devices, San Jose, CA, United States). Endpoint titers were defined by the lowest dilution at which the O.D. was three standard deviations above the mean of the blank wells.

#### SARS-CoV-2 neutralization assays

Microneutralization assays were performed as described previously (Huynh et al., 2021). Briefly, Vero E6 cells (CRL-1586, ATCC, Manassas, VA, United States) were seeded at a density of 2.5 x 10^4^ cells per well in opaque 96 well flat-bottom plates ((Millipore Sigma, Etobicoke, ON, Canada) in complete DMEM (supplemented with 10 % FBS, 1% L-glutamine, 100 U/mL penicillin-streptomycin). 24 hours later, serum was inactivated by incubating at 56°C for 30 minutes, then diluted 1:10 in low serum DMEM (supplemented with 2 % FBS, 1% L-glutamine, 100 U/mL penicillin-streptomycin), followed by a 1:2 dilution series in 96 well U-bottom plates resulting in a final volume of 55 µL diluted serum per well. An equal volume of SARS-CoV-2 consisting of 330 PFU per well was then added to the diluted serum. The serum-virus mixture was then incubated at 37°C for 1 hour. The Vero E6 culture media was then replaced with 100 µL of the serum-virus mixture and was incubated at 37°C for 72 hours. The plates were read by removing 50 µL of culture supernatant and adding 50µL of CellTiter-Glo 2.0 Reagent (Promega, Madison, WI, United States) to each well. The plates were then shaken at 282 cpm at 3 mm diameter for 2 minutes, incubated for 5 minutes at room temperature, then luminescence was read using a BioTek Synergy H1 microplate reader with gain of 135 and integration time of 1 second.

In certain experiments, serum neutralizing antibodies were assessed utilizing a surrogate SARS-CoV-2 virus neutralization test (sVNT). sVNT assays were performed utilizing the cPass Neutralization Antibody Detection kit (GenScript, Piscataway, NJ, United States), according to manufacturer’s instructions.

#### Cytokine and serum chemistry

Evaluation of serum and BAL cytokines were performed by Eve Technologies (Alberta, Canada) through a mouse cytokine array/chemokine array 44-plex (MD44). Serum chemistry was performed by Antech Diagnostics (Ontario, Canada) through a chemistry panel (BioChem 5 panel).

#### Flow cytometry

Cell immunostaining and flow cytometry were performed as previously described (D’Agostino et al., 2020; Jeyanathan et al., 2015; Yao et al., 2018). Briefly, tissue isolated mononuclear cells were plated in U-bottom, 96-well plates at a concentration of 2×10^7^ cells/mL in PBS. Following staining with The LIVE/DEAD™ Fixable Aqua Dead Cell Stain Kit (ThermoFisher Scientific Waltham, MA, United States) at room temperature for 30 min, cells were washed and blocked with anti-CD16/CD32 (clone 2.4G2) in 0.5 % BSA-PBS for 15 min on ice and then stained with fluorochrome-labeled mAbs for 30 min on ice. Fluorochrome-labeled mAbs used for staining cells were anti-CD45–APC-Cy7 (clone 30-F11), anti-CD11b–PE-Cy7 (clone M1/70), anti-CD11c–APC (clone HL3), anti–MHC class II (MHC II)–Alexa Fluor 700 (clone M5/114.15.2; eBioscience, ThermoFisher Scientific Waltham, MA, United States), anti-CD3-V450 (clone 17A2), anti-CD45R (B220)-V450 (clone RA3-6B2), anti–Ly-6C–Biotin (clone HK1.4; BioLegend, San Diego, CA, United States), Streptavidin Qdot 800 (ThermoFisher Scientific Waltham, MA, United States), anti-CD24–BV650 (clone M1/69), anti-CD64–PE (clone 354-5/7.1; BioLegend, San Diego, CA, United States), anti–Ly-6G–BV605 (clone 1A8), anti–Siglec-F–PE-CF594 (clone E50-2440), anti-CD4 APC-Cy7 (clone GK1.5), anti-CD8 PE-Cy7 (clone 53-6.7), anti-IFNγ APC (clone XMG1.2), anti-TNFα FITC (clone MP6-XT22), anti-IL2 BV605 (clone JES6-5H4), anti-Granzyme B PE (clone NGZB; ThermoFisher Scientific Waltham, MA, United States), anti-CD44 PE (clone IM7), anti-CD69 BV605 (clone H1.2F3), anti-CD103-Biotin (clone M290), anti-CD11a FITC (clone 2D7), anti-IgD BV711 (clone 11-26c.2a; BioLegend San Diego, CA, United States), and anti-IgG1-BV421 (clone RMG1-1; BioLegend San Diego, CA, United States). Stained cells were fixed and permeabilized with BD Cytofix/Cytoperm before incubation in BD Perm/Wash buffer (BD Biosciences, San Jose, CA, United States). All mAbs and reagents were purchased from BD Biosciences unless otherwise indicated. Stained cells were processed according to BD Biosciences instructions for flow cytometry and run on a BD LSR II flow cytometer. Data were analyzed using FlowJo software (version 10.1; Tree Star, Ashland, OR, United States).

#### SARS-CoV-2 viral burden determination in tissues

Lung and brains were homogenized using a Bead Mill 24 homogenizer (ThermoFisher Scientific Waltham, MA, United States). Homogenates were clarified by centrifugation at 300 x g and frozen at −80°C. Homogenates were then thawed, and serially diluted 1:10 in serum-free DMEM supplemented with 1% HEPES pH 7.3, 1 mM sodium pyruvate, 1% L-Glutamine and 100 U/mL of penicillin–streptomycin. 100 µL of viral inoculum was plated on Vero E6 cells in 96-well plates (4 x 10^4^ cells per well) for 1hr at 37°C. 5% CO_2_, at which point the inoculum was replaced with low-serum DMEM supplemented with 2% fetal bovine serum (FBS), 1% HEPES pH 7.3, 1 mM sodium pyruvate, 1% L-Glutamine and 100 U/mL of penicillin–streptomycin. Wells were assessed for cytopathic effect at 5-days post-infection using an EVOS M5000 microscope (ThermoFisher Scientific Waltham, MA, United States).

#### Peptide library construction and stimulation

Peptide libraries consisting of 10 amino acid, 15mer synthetic overlapping peptides for vaccine encoded antigens S1, nucleocapsid, and RdRp were synthesized by Pepscan (Lelystad, The Netherlands). Peptides were reconstituted in DMSO according to manufacturer’s instructions to a final concentration of 40 µg/µL. Antigen peptide pools were generated with each pool containing 0.2 µg/µL of each peptide. Unless otherwise stated, peptide stimulations were carried for each vaccine antigen individually with their respective peptide pools, utilizing 2 µg of each peptide/mL of culture media.

### QUANTIFICATION AND STATISTICAL ANALYSIS

Asterisks in the figures indicate the level of statistical significance (*P ≤ 0.05, **P ≤ 0.01, ***P ≤ 0.001, and ****P ≤ 0.0001) as determined using a two-tailed unpaired Student t test, one-way ANOVA, or two-way ANOVA with a Tukey post hoc analysis. Tests were performed using GraphPad Prism software (Version 9, Graphpad Software, La Jolla, CA, United States). Data are expressed as mean ± SEM unless otherwise stated.

## References

Abu-Raddad, L.J., Chemaitelly, H., and Butt, A.A. (2021). Effectiveness of the BNT162b2 Covid-19 Vaccine against the B.1.1.7 and B.1.351 Variants. N. Engl. J. Med. NEJMc2104974.

Alter, G., Yu, J., Liu, J., Chandrashekar, A., Borducchi, E.N., Tostanoski, L.H., McMahan, K., Jacob-Dolan, C., Martinez, D.R., Chang, A., et al. (2021). Immunogenicity of Ad26.COV2.S vaccine against SARS-CoV-2 variants in humans. Nature.

Altmann, D.M., and Boyton, R.J. (2020). SARS-CoV-2 T cell immunity: Specificity, function, durability, and role in protection. Sci. Immunol. 5, eabd6160.

Amanat, F., Stadlbauer, D., Strohmeier, S., Nguyen, T.H.O., Chromikova, V., McMahon, M., Jiang, K., Arunkumar, G.A., Jurczyszak, D., Polanco, J., et al. (2020). A serological assay to detect SARS-CoV-2 seroconversion in humans. Nat. Med. 26, 1033–1036.

Andreano, E., and Rappuoli, R. (2021). SARS-CoV-2 escaped natural immunity, raising questions about vaccines and therapies. Nat. Med. 27, 759–761.

Aschwanden, C. (2021). Five reasons why COVID herd immunity is probably impossible. Nature 591, 520–522.

Banerjee, A., Nasir, J.A., Budylowski, P., Yip, L., Aftanas, P., Christie, N., Ghalami, A., Baid, K., Raphenya, A.R., Hirota, J.A., et al. (2020). Isolation, Sequence, Infectivity, and Replication Kinetics of Severe Acute Respiratory Syndrome Coronavirus 2. Emerg. Infect. Dis. 26, 2054–2063.

Belyakov, I.M., and Ahlers, J.D. (2009). What Role Does the Route of Immunization Play in the Generation of Protective Immunity against Mucosal Pathogens? J. Immunol. 183, 6883–6892.

Bliss, C.M., Parsons, A.J., Nachbagauer, R., Hamilton, J.R., Cappuccini, F., Ulaszewska, M., Webber, J.P., Clayton, A., Hill, A.V.S., and Coughlan, L. (2020). Targeting Antigen to the Surface of EVs Improves the In Vivo Immunogenicity of Human and Non-human Adenoviral Vaccines in Mice. Mol. Ther. - Methods Clin. Dev. 16, 108–125.

Bournazos, S., Gupta, A., and Ravetch, J. V. (2020). The role of IgG Fc receptors in antibody-dependent enhancement. Nat. Rev. Immunol. 20, 633–643.

Bricker, T.L., Darling, T.L., Hassan, A.O., Harastani, H.H., Soung, A., Jiang, X., Dai, Y.-N., Zhao, H., Adams, L.J., Holtzman, M.J., et al. (2021). A single intranasal or intramuscular immunization with chimpanzee adenovirus vectored SARS-CoV-2 vaccine protects against pneumonia in hamsters. Cell Rep. 109400.

Callaway, E., and Ledford, H. (2021). How to redesign COVID vaccines so they protect against variants. Nature 590, 15–16.

Cao, H., Mai, J.M., Zhou, Z., Li, Z., Duan, R., Watt, J., Chen, Z., Bandara, R.A., Li, M., Ahn, S.K., et al. (2021). Intranasal HD-Ad Vaccine Protects the Upper and Lower Respiratory Tracts of hACE2 Mice against SARS-CoV-2. BioRxiv 2021.04.08.439006.

Chen, R.E., Zhang, X., Case, J.B., Winkler, E.S., Liu, Y., VanBlargan, L.A., Liu, J., Errico, J.M., Xie, X., Suryadevara, N., et al. (2021). Resistance of SARS-CoV-2 variants to neutralization by monoclonal and serum-derived polyclonal antibodies. Nat. Med. 27, 717–726.

Class, J., Dangi, T., Richner, J.M., Penaloza-Macmaster, P., and Richner, J. (2021). A SARS CoV-2 nucleocapsid vaccine protects against distal viral dissemination. BioRxiv 2021.04.26.440920.

D’Agostino, M.R., Lai, R., Afkhami, S., Khera, A., Yao, Y., Vaseghi-Shanjani, M., Zganiacz, A., Jeyanathan, M., and Xing, Z. (2020). Airway Macrophages Mediate Mucosal Vaccine–Induced Trained Innate Immunity against Mycobacterium tuberculosis in Early Stages of Infection. J. Immunol. 205, 2750–2762.

Dai, L., and Gao, G.F. (2021). Viral targets for vaccines against COVID-19. Nat. Rev. Immunol. 21, 73–82.

Fontanet, A., and Cauchemez, S. (2020). COVID-19 herd immunity: where are we? Nat. Rev. Immunol. 20, 583–584.

Garcia-Beltran, W.F., Lam, E.C., St. Denis, K., Nitido, A.D., Garcia, Z.H., Hauser, B.M., Feldman, J., Pavlovic, M.N., Gregory, D.J., Poznansky, M.C., et al. (2021). Multiple SARS-CoV-2 variants escape neutralization by vaccine-induced humoral immunity. Cell 184, 2372–2383.e9.

Geers, D., Shamier, M.C., Bogers, S., den Hartog, G., Gommers, L., Nieuwkoop, N.N., Schmitz, K.S., Rijsbergen, L.C., van Osch, J.A.T., Dijkhuizen, E., et al. (2021). SARS-CoV-2 variants of concern partially escape humoral but not T-cell responses in COVID-19 convalescent donors and vaccinees. Sci. Immunol. 6, eabj1750.

Grant, R.A., Morales-Nebreda, L., Markov, N.S., Swaminathan, S., Querrey, M., Guzman, E.R., Abbott, D.A., Donnelly, H.K., Donayre, A., Goldberg, I.A., et al. (2021). Circuits between infected macrophages and T cells in SARS-CoV-2 pneumonia. Nature 590, 635–641.

Gupta, R.K. (2021). Will SARS-CoV-2 variants of concern affect the promise of vaccines? Nat. Rev. Immunol. 21, 340–341.

Hartley, G.E., Edwards, E.S.J., Aui, P.M., Varese, N., Stojanovic, S., McMahon, J., Peleg, A.Y., Boo, I., Drummer, H.E., Hogarth, P.M., et al. (2020). Rapid generation of durable B cell memory to SARS-CoV-2 spike and nucleocapsid proteins in COVID-19 and convalescence. Sci. Immunol. 5, eabf8891.

Harvey, W.T., Carabelli, A.M., Jackson, B., Gupta, R.K., Thomson, E.C., Harrison, E.M., Ludden, C., Reeve, R., Rambaut, A., Peacock, S.J., et al. (2021). SARS-CoV-2 variants, spike mutations and immune escape. Nat. Rev. Microbiol. 19, 409–424.

Hassan, A.O., Kafai, N.M., Dmitriev, I.P., Fox, J.M., Smith, B.K., Harvey, I.B., Chen, R.E., Winkler, E.S., Wessel, A.W., Case, J.B., et al. (2020). A Single-Dose Intranasal ChAd Vaccine Protects Upper and Lower Respiratory Tracts against SARS-CoV-2. Cell 183, 169–184.e13.

Hassan, A.O., Feldmann, F., Zhao, H., Curiel, D.T., Okumura, A., Tang-Huau, T.-L., Case, J.B., Meade-White, K., Callison, J., Chen, R.E., et al. (2021a). A single intranasal dose of chimpanzee adenovirus-vectored vaccine protects against SARS-CoV-2 infection in rhesus macaques. Cell Reports Med. 2, 100230.

Hassan, A.O., Shrihari, S., Gorman, M.J., Ying, B., Yuan, D., Raju, S., Chen, R.E., Dmitriev, I.P., Kashentseva, E., Adams, L.J., et al. (2021b). An intranasal vaccine durably protects against SARS-CoV-2 variants in mice. BioRxiv 2021.05.08.443267.

Hoffmann, D., Corleis, B., Rauch, S., Roth, N., Mühe, J., Halwe, N.J., Ulrich, L., Fricke, C., Schön, J., Kraft, A., et al. (2021a). CVnCoV and CV2CoV protect human ACE2 transgenic mice from ancestral B BavPat1 and emerging B.1.351 SARS-CoV-2. Nat. Commun. 12, 4048.

Hoffmann, M., Arora, P., Groß, R., Seidel, A., Hörnich, B.F., Hahn, A.S., Krüger, N., Graichen, L., Hofmann-Winkler, H., Kempf, A., et al. (2021b). SARS-CoV-2 variants B.1.351 and P.1 escape from neutralizing antibodies. Cell 184, 2384–2393.e12.

Huang, A.T., Garcia-Carreras, B., Hitchings, M.D.T., Yang, B., Katzelnick, L.C., Rattigan, S.M., Borgert, B.A., Moreno, C.A., Solomon, B.D., Trimmer-Smith, L., et al. (2020). A systematic review of antibody mediated immunity to coronaviruses: kinetics, correlates of protection, and association with severity. Nat. Commun. 11, 4704.

Huynh, A., Arnold, D.M., Smith, J.W., Moore, J.C., Zhang, A., Chagla, Z., Harvey, B.J., Stacey, H.D., Ang, J.C., Clare, R., et al. (2021). Characteristics of Anti-SARS-CoV-2 Antibodies in Recovered COVID-19 Subjects. Viruses 13, 697.

Jeyanathan, M., Thanthrige-Don, N., Afkhami, S., Lai, R., Damjanovic, D., Zganiacz, A., Feng, X., Yao, X.-D., Rosenthal, K.L., Fe Medina, M., et al. (2015). Novel chimpanzee adenovirus-vectored respiratory mucosal tuberculosis vaccine: overcoming local anti-human adenovirus immunity for potent TB protection. Mucosal Immunol. 8, 1373–1387.

Jeyanathan, M., Afkhami, S., Khera, A., Mandur, T., Damjanovic, D., Yao, Y., Lai, R., Haddadi, S., Dvorkin-Gheva, A., Jordana, M., et al. (2017). CXCR3 Signaling Is Required for Restricted Homing of Parenteral Tuberculosis Vaccine–Induced T Cells to Both the Lung Parenchyma and Airway. J. Immunol. 199, 2555–2569.

Jeyanathan, M., Yao, Y., Afkhami, S., Smaill, F., and Xing, Z. (2018). New Tuberculosis Vaccine Strategies: Taking Aim at Un-Natural Immunity. Trends Immunol. 39, 419–433.

Jeyanathan, M., Afkhami, S., Smaill, F., Miller, M.S., Lichty, B.D., and Xing, Z. (2020). Immunological considerations for COVID-19 vaccine strategies. Nat. Rev. Immunol. 20, 615–632.

Khoury, D.S., Wheatley, A.K., Ramuta, M.D., Reynaldi, A., Cromer, D., Subbarao, K., O’Connor, D.H., Kent, S.J., and Davenport, M.P. (2020). Measuring immunity to SARS-CoV-2 infection: comparing assays and animal models. Nat. Rev. Immunol. 20, 727–738.

Khoury, D.S., Cromer, D., Reynaldi, A., Schlub, T.E., Wheatley, A.K., Juno, J.A., Subbarao, K., Kent, S.J., Triccas, J.A., and Davenport, M.P. (2021). Neutralizing antibody levels are highly predictive of immune protection from symptomatic SARS-CoV-2 infection. Nat. Med.

Krammer, F. (2021). Correlates of protection from SARS-CoV-2 infection. Lancet 397, 1421–1423.

Krause, P.R., Fleming, T.R., Longini, I.M., Peto, R., Briand, S., Heymann, D.L., Beral, V., Snape, M.D., Rees, H., Ropero, A.-M., et al. (2021). SARS-CoV-2 Variants and Vaccines. N. Engl. J. Med. NEJMsr2105280.

Ku, M.-W., Bourgine, M., Authié, P., Lopez, J., Nemirov, K., Moncoq, F., Noirat, A., Vesin, B., Nevo, F., Blanc, C., et al. (2021). Intranasal vaccination with a lentiviral vector protects against SARS-CoV-2 in preclinical animal models. Cell Host Microbe 29, 236–249.e6.

Kuate, S., Cinatl, J., Doerr, H.W., and Überla, K. (2007). Exosomal vaccines containing the S protein of the SARS coronavirus induce high levels of neutralizing antibodies. Virology 362, 26–37.

Kumagai, Y., Takeuchi, O., Kato, H., Kumar, H., Matsui, K., Morii, E., Aozasa, K., Kawai, T., and Akira, S. (2007). Alveolar Macrophages Are the Primary Interferon-α Producer in Pulmonary Infection with RNA Viruses. Immunity 27, 240–252.

Lai, R., Afkhami, S., Haddadi, S., Jeyanathan, M., and Xing, Z. (2015). Mucosal immunity and novel tuberculosis vaccine strategies: route of immunisation-determined T-cell homing to restricted lung mucosal compartments. Eur. Respir. Rev. 24, 356–360.

Leist, S.R., Dinnon, K.H., Schäfer, A., Tse, L. V., Okuda, K., Hou, Y.J., West, A., Edwards, C.E., Sanders, W., Fritch, E.J., et al. (2020). A Mouse-Adapted SARS-CoV-2 Induces Acute Lung Injury and Mortality in Standard Laboratory Mice. Cell 183, 1070–1085.e12.

Lurie, N., Saville, M., Hatchett, R., and Halton, J. (2020). Developing Covid-19 Vaccines at Pandemic Speed. N. Engl. J. Med. 382, 1969–1973.

Madhi, S.A., Baillie, V., Cutland, C.L., Voysey, M., Koen, A.L., Fairlie, L., Padayachee, S.D., Dheda, K., Barnabas, S.L., Bhorat, Q.E., et al. (2021). Efficacy of the ChAdOx1 nCoV-19 Covid-19 Vaccine against the B.1.351 Variant. N. Engl. J. Med. 384, 1885–1898.

Matchett, W.E., Joag, V., Stolley, J.M., Shepherd, F.K., Quarnstrom, C.F., Mickelson, C.K., Wijeyesinghe, S., Soerens, A.G., Becker, S., Thiede, J.M., et al. (2021). Nucleocapsid Vaccine Elicits Spike-Independent SARS-CoV-2 Protective Immunity. J. Immunol. ji2100421.

Neutra, M.R., and Kozlowski, P.A. (2006). Mucosal vaccines: the promise and the challenge. Nat. Rev. Immunol. 6, 148–158.

Peng, Y., Mentzer, A.J., Liu, G., Yao, X., Yin, Z., Dong, D., Dejnirattisai, W., Rostron, T., Supasa, P., Liu, C., et al. (2020). Broad and strong memory CD4+ and CD8+ T cells induced by SARS-CoV-2 in UK convalescent individuals following COVID-19. Nat. Immunol. 21, 1336–1345.

Planas, D., Bruel, T., Grzelak, L., Guivel-Benhassine, F., Staropoli, I., Porrot, F., Planchais, C., Buchrieser, J., Rajah, M.M., Bishop, E., et al. (2021). Sensitivity of infectious SARS-CoV-2 B.1.1.7 and B.1.351 variants to neutralizing antibodies. Nat. Med. 27, 917–924.

Rodda, L.B., Netland, J., Shehata, L., Pruner, K.B., Morawski, P.A., Thouvenel, C.D., Takehara, K.K., Eggenberger, J., Hemann, E.A., Waterman, H.R., et al. (2021). Functional SARS-CoV-2-Specific Immune Memory Persists after Mild COVID-19. Cell 184, 169–183.e17.

Sadoff, J., Gray, G., Vandebosch, A., Cárdenas, V., Shukarev, G., Grinsztejn, B., Goepfert, P.A., Truyers, C., Fennema, H., Spiessens, B., et al. (2021). Safety and Efficacy of Single-Dose Ad26.COV2.S Vaccine against Covid-19. N. Engl. J. Med. 384, 2187–2201.

Santosuosso, M., Zhang, X., McCormick, S., Wang, J., Hitt, M., and Xing, Z. (2005). Mechanisms of Mucosal and Parenteral Tuberculosis Vaccinations: Adenoviral-Based Mucosal Immunization Preferentially Elicits Sustained Accumulation of Immune Protective CD4 and CD8 T Cells within the Airway Lumen. J. Immunol. 174, 7986–7994.

Satti, I., Meyer, J., Harris, S.A., Thomas, Z.-R.M., Griffiths, K., Antrobus, R.D., Rowland, R., Ramon, R.L., Smith, M., Sheehan, S., et al. (2014). Safety and immunogenicity of a candidate tuberculosis vaccine MVA85A delivered by aerosol in BCG-vaccinated healthy adults: a phase 1, double-blind, randomised controlled trial. Lancet Infect. Dis. 14, 939–946.

Schneider, C., Nobs, S.P., Heer, A.K., Kurrer, M., Klinke, G., van Rooijen, N., Vogel, J., and Kopf, M. (2014). Alveolar Macrophages Are Essential for Protection from Respiratory Failure and Associated Morbidity following Influenza Virus Infection. PLoS Pathog. 10, e1004053.

Shen, X., Tang, H., Pajon, R., Smith, G., Glenn, G.M., Shi, W., Korber, B., and Montefiori, D.C. (2021). Neutralization of SARS-CoV-2 Variants B.1.429 and B.1.351. N. Engl. J. Med. 384, 2352–2354.

Shinde, V., Bhikha, S., Hoosain, Z., Archary, M., Bhorat, Q., Fairlie, L., Lalloo, U., Masilela, M.S.L., Moodley, D., Hanley, S., et al. (2021). Efficacy of NVX-CoV2373 Covid-19 Vaccine against the B.1.351 Variant. N. Engl. J. Med. 384, 1899–1909.

Singh, R., Kang, A., Luo, X., Jeyanathan, M., Gillgrass, A., Afkhami, S., and Xing, Z. (2021). COVID-19: Current knowledge in clinical features, immunological responses, and vaccine development. FASEB J. 35, 1–23.

Stacey, H.D., Golubeva, D., Posca, A., Ang, J.C., Novakowski, K.E., Zahoor, M.A., Kaushic, C., Cairns, E., Bowdish, D.M.E., Mullarkey, C.E., et al. (2021). IgA potentiates NETosis in response to viral infection. Proc. Natl. Acad. Sci. 118, e2101497118.

Stadlbauer, D., Tan, J., Jiang, K., Hernandez, M.M., Fabre, S., Amanat, F., Teo, C., Arunkumar, G.A., McMahon, M., Capuano, C., et al. (2021). Repeated cross-sectional sero-monitoring of SARS-CoV-2 in New York City. Nature 590, 146–150.

Starr, T.N., Greaney, A.J., Dingens, A.S., and Bloom, J.D. (2021). Complete map of SARS-CoV-2 RBD mutations that escape the monoclonal antibody LY-CoV555 and its cocktail with LY-CoV016. Cell Reports Med. 2, 100255.

Szabo, P.A., Miron, M., and Farber, D.L. (2019). Location, location, location: Tissue resident memory T cells in mice and humans. Sci. Immunol. 4, eaas9673.

Tan, C.W., Chia, W.N., Qin, X., Liu, P., Chen, M.I.-C., Tiu, C., Hu, Z., Chen, V.C.-W., Young, B.E., Sia, W.R., et al. (2020). A SARS-CoV-2 surrogate virus neutralization test based on antibody-mediated blockage of ACE2–spike protein–protein interaction. Nat. Biotechnol. 38, 1073–1078.

Tarke, A., Sidney, J., Methot, N., Zhang, Y., Dan, J.M., Goodwin, B., Rubiro, P., Sutherland, A., da Silva Antunes, R., Frazier, A., et al. (2021). Negligible impact of SARS-CoV-2 variants on CD4 + and CD8 + T cell reactivity in COVID-19 exposed donors and vaccinees. BioRxiv Prepr. Serv. Biol.

Taylor, J.J., Martinez, R.J., Titcombe, P.J., Barsness, L.O., Thomas, S.R., Zhang, N., Katzman, S.D., Jenkins, M.K., and Mueller, D.L. (2012). Deletion and anergy of polyclonal B cells specific for ubiquitous membrane-bound self-antigen. J. Exp. Med. 209, 2065–2077.

Teijaro, J.R., and Farber, D.L. (2021). COVID-19 vaccines: modes of immune activation and future challenges. Nat. Rev. Immunol. 21, 195–197.

Wang, J., Thorson, L., Stokes, R.W., Santosuosso, M., Huygen, K., Zganiacz, A., Hitt, M., and Xing, Z. (2004). Single mucosal, but not parenteral, immunization with recombinant adenoviral-based vaccine provides potent protection from pulmonary tuberculosis. J. Immunol. 173, 6357–6365.

Wang, P., Nair, M.S., Liu, L., Iketani, S., Luo, Y., Guo, Y., Wang, M., Yu, J., Zhang, B., Kwong, P.D., et al. (2021). Antibody resistance of SARS-CoV-2 variants B.1.351 and B.1.1.7. Nature 593, 130–135.

Xing, Z., Afkhami, S., Bavananthasivam, J., Fritz, D.K., D’Agostino, M.R., Vaseghi-Shanjani, M., Yao, Y., and Jeyanathan, M. (2020). Innate immune memory of tissue-resident macrophages and trained innate immunity: Re-vamping vaccine concept and strategies. J. Leukoc. Biol. 108, 825–834.

Yao, Y., Jeyanathan, M., Haddadi, S., Barra, N.G., Vaseghi-Shanjani, M., Damjanovic, D., Lai, R., Afkhami, S., Chen, Y., Dvorkin-Gheva, A., et al. (2018). Induction of Autonomous Memory Alveolar Macrophages Requires T Cell Help and Is Critical to Trained Immunity. Cell 175, 1634–1650.e17.

Zhao, J., Zhao, J., Mangalam, A.K., Channappanavar, R., Fett, C., Meyerholz, D.K., Agnihothram, S., Baric, R.S., David, C.S., and Perlman, S. (2016). Airway Memory CD4 + T Cells Mediate Protective Immunity against Emerging Respiratory Coronaviruses. Immunity 44, 1379–1391.

Zheng, J., Wong, L.Y.R., Li, K., Verma, A.K., Ortiz, M.E., Wohlford-Lenane, C., Leidinger, M.R., Knudson, C.M., Meyerholz, D.K., McCray, P.B., et al. (2021). COVID-19 treatments and pathogenesis including anosmia in K18-hACE2 mice. Nature 589, 603–607.

